# A G protein-coupled receptor is required in cartilaginous and dense connective tissues to maintain spine alignment

**DOI:** 10.1101/2021.02.22.432353

**Authors:** Zhaoyang Liu, Amro A. Hussien, Yunjia Wang, Terry Heckmann, Roberto Gonzalez, Courtney M. Karner, Jess G. Snedeker, Ryan S. Gray

## Abstract

Adolescent idiopathic scoliosis (AIS) is the most common spine disorder affecting children worldwide, yet little is known about the pathogenesis of this disorder. Here, we demonstrate that genetic regulation of structural components of the axial skeleton, the intervertebral discs and dense connective tissues (e.g., ligaments and tendons), are essential for maintenance of spinal alignment. We show that the G-coupled protein receptor *Adgrg6*, previously implicated in human AIS association studies, is required in these tissues to maintain typical spine morphology. We show that *Adgrg6* regulates biomechanical properties of tendon and stimulates CREB signaling governing gene expression in cartilaginous tissues of the spine. Treatment with an cAMP agonist was able to mirror aspects of receptor function in culture defining core pathways for regulation of these axial connective tissues. As *ADGRG6* is a key gene involved in human AIS, these findings open up novel therapeutic opportunities for human scoliosis.

**Highlights:**

- Knockout mice lacking *Adgrg6* function in the tendons and ligaments of the spine develop perinatal-onset thoracic scoliosis.
- Loss of *Adgrg6* function in cartilaginous tissues of the discs contribute to the incidence and severity of scoliosis.
- The loss of *Adgrg6* function in spine tissues provide a model of construct validity for human adolescent idiopathic scoliosis
- Fine tuning of the biomechanical properties of dense connective tissues is essential for maintaining spine alignment.

## Introduction

The maturation and homeostasis of a healthy, functional spine requires the integration of several musculoskeletal tissues including: bone, cartilage and connective tissues, muscle, and the peripheral nervous system. The spine consists of a series of segmented bony vertebral bodies which are linked together by fibrocartilaginous joints named the intervertebral discs (IVDs), which aid in lateral and rotational flexibility and cushioning for axial loading of the spine. The spine is further supported by the paraspinal ligaments and provides attachment sites for multiple muscle-tendon insertions. Finally, the structural organization of the axial spine aids in the protection of the spinal cord and acts as the central axis of the body to support and maintain posture during movement through the environment (Bagnat and Gray, 2020; Bogduk, 2016).

Scoliosis is a complex rotational atypical configuration of the spine, which is commonly diagnosed by a lateral curvature of the spine of 10° or greater in the coronal plane (Cheng et al., 2015). Scoliosis can be caused by congenital patterning defects of the vertebral column or associated with a variety of neuromuscular or syndromic disorders (Murphy and Mooney, 2019). Whereas, idiopathic scoliosis occurs in otherwise healthy children without associated patterning or neuro-muscular conditions. Adolescent idiopathic scoliosis (AIS) is the most common pediatric spine disorder, affecting ∼3% of children worldwide (Cheng et al., 2015). Clinical interventions for AIS includes surgical correction which aim to halt the progression of severe curves, but which places a high socio-economic burden on patients and families (Negrini et al., 2018). Despite substantial efforts to decipher the pathogenesis of AIS, the molecular genetics and pathology underlying this condition remain ill-defined (Cheng et al., 2015; Newton Ede and Jones, 2016).

The appearance of spine curvatures in AIS patients without obvious morphological defects of the vertebral column suggests that more subtle defects in one or more regulators of spine stability may contribute to pathogenesis of this disorder. Indeed, subtle asymmetric morphological changes of the vertebral bodies at the apical region of the curve have been observed in some AIS patients, which may initiate spine curvature (Lam et al., 2011; Liljenqvist et al., 2002). In addition, changes of cell density and glycosaminoglycan composition (Shu and Melrose, 2018; Urban et al., 2001); altered ultrastructure of collagen and elastic fibers (Akhtar et al., 2005); and increased incidence of endplate-oriented disc herniations (e.g. Schmorl’s nodes) (Buttermann and Mullin, 2008) may also contribute to the onset of scoliosis in AIS patients.

Genome-wide association studies and subsequent meta-analysis of multi-ethnic AIS cohorts strongly implicate the *ADGRG6* gene for susceptibility to scoliosis (Kou et al., 2013; Kou et al., 2018). *Adgrg6* (also called *Gpr126*) is a member of the adhesion G protein-coupled receptor family of proteins, most of which display canonical intercellular signaling function via activation of cyclic adenosine monophosphate (cAMP) (Hamann et al., 2015; Langenhan et al., 2016). ADGRG6 has been shown to function through canonical cAMP signaling for the regulation of Schwann cell differentiation and myelination, as well as, in regulation of inner ear development (Geng et al., 2013; Monk et al., 2009; Monk et al., 2011). *Adgrg6* null mutant mice are embryonic lethal, with a few survivors displaying cardiac defects and severe joint contractures, but which die before weaning (Mogha et al., 2013; Monk et al., 2011; Waller-Evans et al., 2010). However, conditional ablation of *Adgrg6* in osteochondral progenitor cells generated a reproducible genetic mouse model which displayed postnatal-onset scoliosis, without generating obvious patterning defects of the vertebral column (Karner et al., 2015). It remains to be determined, however, which elements of the spine and signaling pathways are important for maintaining normal spine alignment and how alteration of these deficiencies may alleviate pathology and distress in patients.

In this work, we identify a unique target of *Adgrg6* signaling whereby the ligaments and tendons and intervertebral discs act synergistically to maintain spine alignment, rather than bony tissues. We define signaling mechanisms of *Adgrg6* that provide foundations to advance the understanding of the pathogenesis of AIS and provide a mouse models of AIS with a high degree on construct validity (Willner, 1984) for furthering pharmacological-based interventions of this disorder.

## Results

### Ablation of *Adgrg6* in osteochondral progenitor cells models progressive adolescent idiopathic scoliosis in mouse

Multi-ethnic genome wide association studies show that intronic variants *GPR126/ADGRG6* locus are associated with AIS in human (Kou et al., 2013; Kou et al., 2018). Conditional knockout (cKO) of *Adgrg6* in osteochondral progenitor cells (*Col2Cre;Adgrg6^f/f^*, called *Col2-cKO* for short) models the timing and pathology of AIS in mouse (Karner et al., 2015), which makes this mouse model uniquely suited to study the initiation, progression, and pathogenesis of AIS. To determine the natural history of scoliosis in *Col2-cKO* mice, we performed longitudinal X-ray analysis at postnatal day 10 (P10) out to P120. Both wild-type (WT) Cre (-) and heterozygous Cre (+) (*Col2Cre; Adgrg6^f/+^*) control littermates showed typical patterning of a straight spinal column at all timepoints (Fig. 1A-D’ and Fig. S1A, A’). *Col2-cKO* mice were indistinguishable from littermate controls at birth and displayed typical patterning and alignment of the spine at P10 (0/23) (Fig. 1E, E’). However, at P20 we first observed postnatal-onset spine curvatures (Fig. 1F, F’), which could be progressive in some *Col2-cKO* animals (Fig. 1F-H’). Approximately 50-60% of *Col2-cKO* mice exhibited scoliosis at P20 (12/23), P40 (14/23), and P120 (10/17) (Fig. 1I, K, L), with the apex of the curvature centered at thoracic vertebrae (T)8 and T9 (Fig. 1J). Scoliosis is defined by a Cobb angle measure greater than 10° in the coronal plane (Cobb, 1958), which was increased in *Col2-cKO* mice compared with control littermates (Fig. 1I), ranging from 11° to 46° in scoliotic individuals (Fig. 1I and L). In comparison, only one Cre (+) control mouse (1/18) has a mild curve of 13.8° at P120 and the rest had no measurable alteration in spine alignment (Fig 1I, K and Fig. S1B). We observed scoliosis in both 37.5% (3/8) of male and 66.6% (10/15) of female mutant mice, with both right- or left-ward curvatures (Fig. 1K, L). We did not observe an obvious sex bias for the incidence of scoliosis (Fisher exact test, *p*=0.2213).

**Figure 1.**
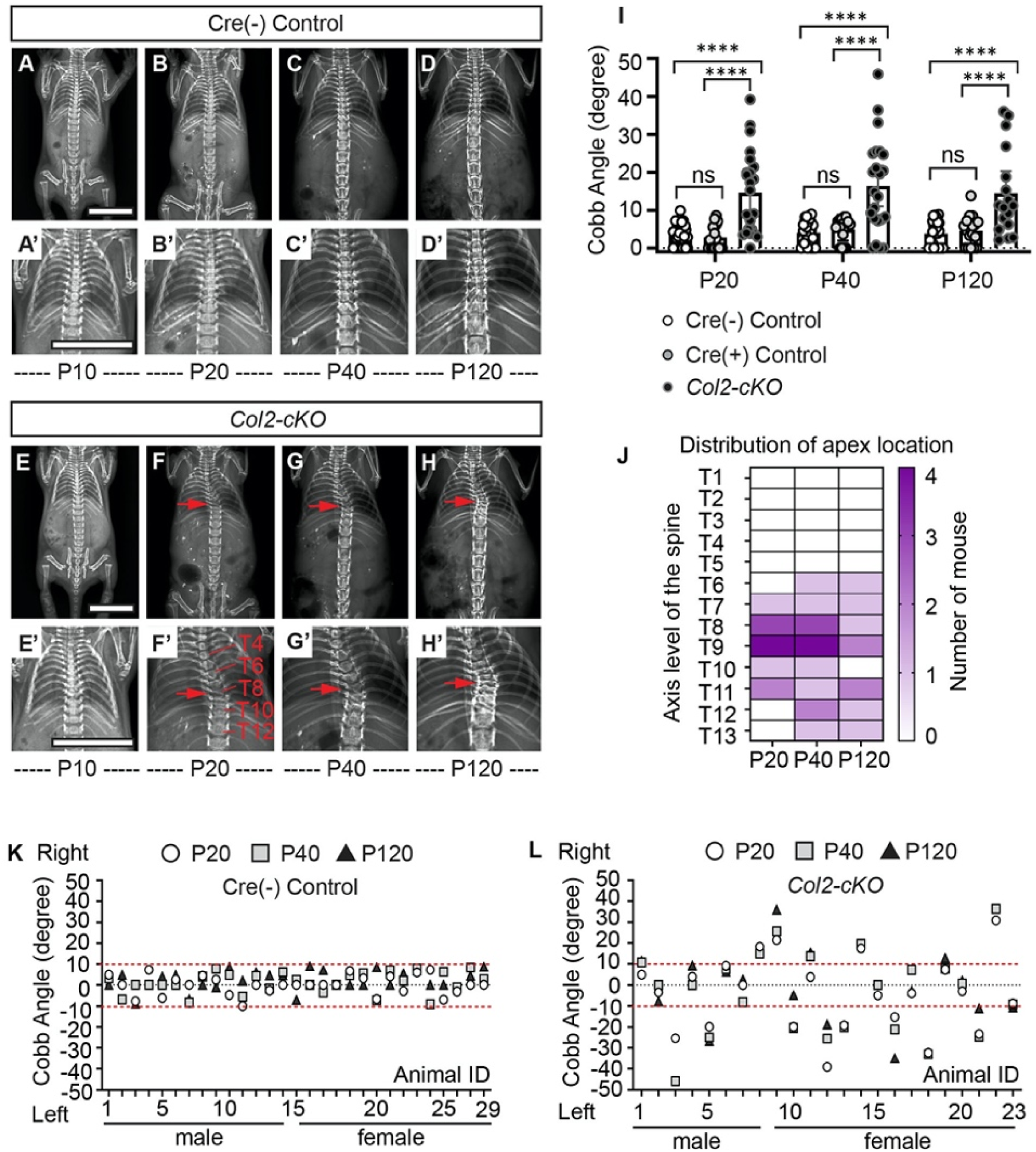
Ablation of *Adgrg6* in osteochondral progenitor cells models progressive AIS in mouse. (A-D’) Longitudinal X-ray analysis of a representative Cre (-) control mouse at P10 (A, A’), P20 (B, B’), P40 (C, C’), and P120 (D, D’). (E-H’) Longitudinal X-ray analysis of a representative *Col2-cKO* mutant mouse at P10 (E, E’), P20 (F, F’), P40 (G, G’), and P120 (H, H’) that shows adolescent-onset (F) and progressive (F-H) thoracic scoliosis. The apexes of scoliosis are indicated with red arrows in (F-H’). Thoracic spine levels are labeled in (F’). (I) Longitudinal Cobb angle analysis of Cre (-) control mice, Cre (+) control mice, and *Col2-cKO* mutant mice at P20, P40 and P120. Cre (-) control mice, *n*=29 mice at P20, P40 and P120; for Cre (+) control mice, *n*=18, 17 and 18 mice at P20, P40, and P120, respectively; for *Col2-cKO* mutant mice, *n*=23, 23 and 17 mice at P20, P40, and P120, respectively. Dots are plotted with mean ± 95% CI. ****: *p*<0.0001, Two-Way ANOVA followed by Tukey test. ns: not significant. (J) Heat map of the apex distribution of the scoliotic *Col2-cKO* mice at P20, P40 and P120. Heat map is plotted with the axis level of the thoracic spine (T1-T13, left axis), and the number of scoliotic mouse (right axis) with apex observed at each level. Apex of scoliosis is distributed alone the middle to lower thoracic spine (T6-T13), with hotspots at T8 and T9. (K, L) Cobb angle values for all the Cre (-) control mice (K) and *Col2-cKO* mice (L) showed in (I). Thresholds of scoliosis (Cobb angle >10°) are indicated with two red dot lines. Scale bars: 10mm.

### Wedging of the intervertebral disc is concurrent with the onset of scoliosis in *Col2-cKO* mutant mice

Wedging of the vertebrae and the IVD within regions of acute spine curvature are commonly observed in AIS (Little et al., 2016; Newton Ede and Jones, 2016). To determine if similar morphological changes of the spine were associated with the initiation and progression of scoliosis, we harvested the thoracic spine (T5-T12) of *Col2-cKO* and control littermates at P20 and P120. During early initiation of scoliosis (P20) we observed wedging of IVDs close to the apex of the curvature in the *Col2-cKO* mice, which was associated with a shift of the nucleus pulposus towards the convexity of the curve (Fig. 2B). The large vacuolated cells (magenta stained/notochordal cells) shifted towards the convex side, while the smaller non-vacuolated cells (blue stained/chondrocyte-like cells) distributed in the middle and concave side (Fig. 2B, B”), in contrast to the uniform distribution of these cells with the IVD typical of control littermates (Fig. 2A, A”). We observed more vertical alignment of the annulus fibrosis lamellae in *Col2-cKO* mice along the convex side (white dash lines, Fig. 2A’ and B’), with flatter lamellae and an elongated inner annulus fibrosus at the concave side the IVD (white dash lines, Fig. 2A and B).

**Figure 2.**
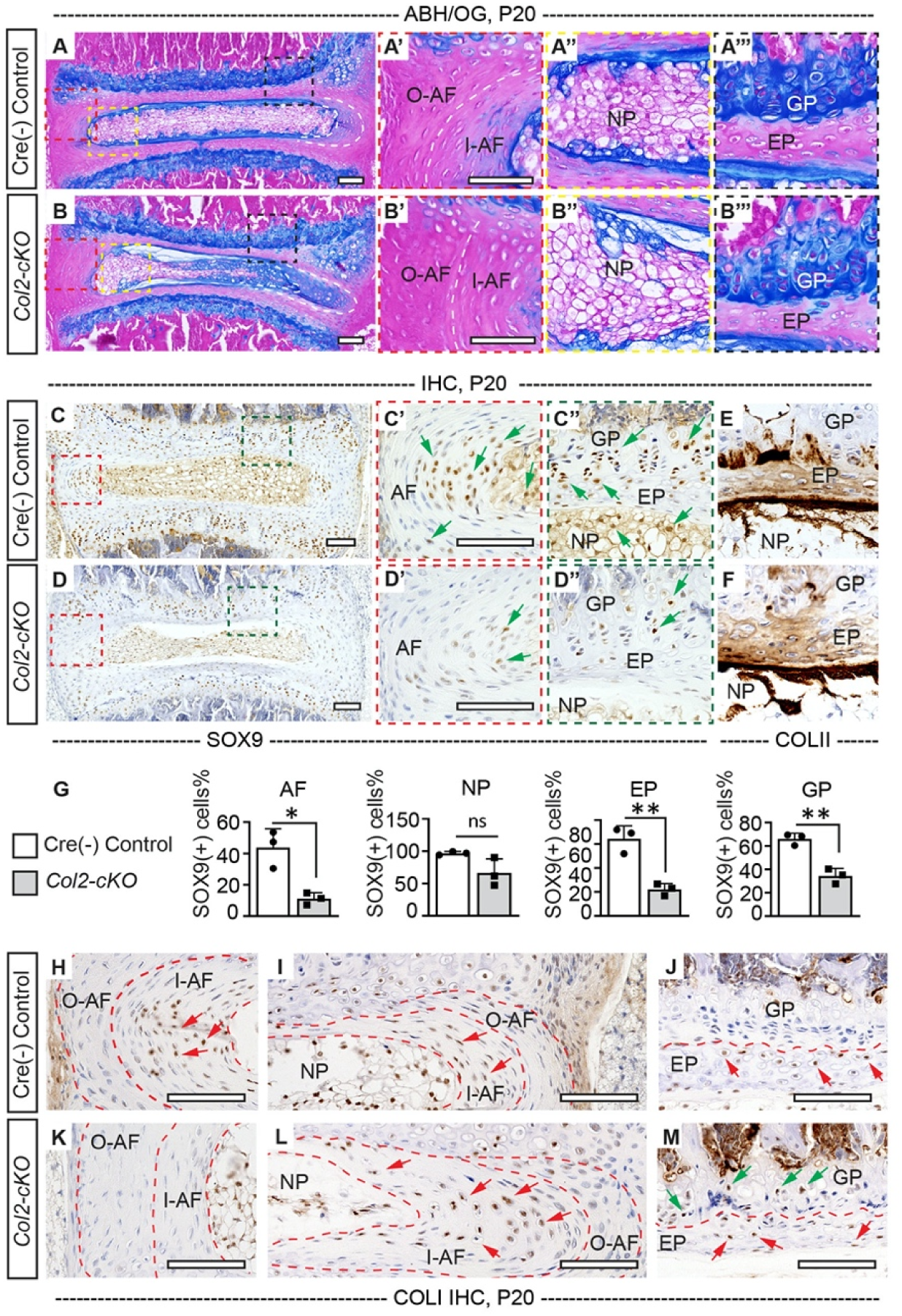
*Col2-cKO* mice display alterations in intervertebral discs. (A-B’’’) Representative intervertebral disc (IVD) sections of Cre (-) control and *Col2-cKO* mutant mice stained with Alcian Blue Hematoxylin/Orange G (ABH/OG) at P20. The IVD close to the apex of the curve in the *Col2-cKO* mice are mildly wedged, associated with a shift in position of the nucleus pulposus towards the convex side of the curve (B, B”). The inner AF (I-AF) and outer AF (O-AF) in the convex side of the *Col2-cKO* mice is composed of more vertical lamellae compared with that in Cre (-) control mice (white dash lines, A’ and B’). The AF in the concave side of the *Col2-cKO* mice is elongated and composed with flatter lamellae, compared with that in Cre (-) control mice (white dash lines, A and B). No overt structural defects are observed in the EP and GP of the *Col2-cKO* mice (B’’’). *n*=4 mice for each group. (C-G) IHC analyses of common anabolic markers of healthy disc at P20. *Col2-cKO* mutant mice display reduced expression of SOX9 in AF, EP, and GP (green arrows, C’-D”), which is quantified in (G). *Col2-cKO* mice show normal expression pattern of COLII compared with the controls (E, F). *n*=3 mice for each group. Bars are plotted with mean and SD. Each dot represents one IVD analyzed. **: *p*<0.01, two-tailed Student’s *t* test. ns: not significant. (H-M) IHC analyses of COLI at P20. COLI is mainly expressed in the I-AF (H, I) and some cells of the EP (J) in the control mice (indicated with red arrows). No obvious expression of COLI is observed in the O-AF or I-AF in the convex side of the mutant IVD (K), but robust expression is observed in the elongate I-AF in the concave side of the mutant IVD (red arrows, L). Notably, some cells in the GP of the *Col2-cKO* mice also express COLI (green arrows, M). *n*=3 mice for each group. Scale bars: 100μm. *AF-annulus fibrosis, I-AF-inner annulus fibrosis, O-AFouter annulus fibrosis, EP- endplate, GP- growth plate, and NP- nucleus pulposus*.

Older *Col2-cKO* scoliotic mice (P120) showed a more pronounced shift of the nucleus pulposus to the convex side (Fig. S1F, F”), with vertical alignment of the annulus fibrosus lamellae (Fig. S1F, F’) and the endplate and growth plate tissues appeared thinner (Fig. S1E’’’, F’’’). Analysis of the cartilaginous endplate and the vertebral growth plate in *Col2-cKO* mice showed no obvious histopathology at P20 (Fig. 2A, A’’’ and B, B’’’). Despite wedging of IVDs close to the apex of the curvature in *Col2-cKO* mice, no apparent defects in the size or patterning of the vertebrae, growth plates, or IVD tissues were observed during the initiation of scoliosis (Fig. S1C, D). However, we did occasionally observe bulging of the cortical bone outward on the concave side of the curve (50%; n=4; black arrow, Fig S1D). Interestingly, the IVDs lying proximal or distal to obvious spine curvatures in *Col2-cKO* mice consistently exhibited typical morphology of the nucleus pulposus and annulus fibrosus comparable with the control littermates (Fig. S1C’, D’), suggesting that localized forces at the thoracic spine are driving the alterations in tissue architecture.

### *Col2-cKO* mutant mice show a reduction in SOX9 protein expression without significant alterations of IVD matrix composition at the onset of scoliosis

Analysis of the IVD and growth plate in human AIS patients have demonstrated alterations in cellularity, changes in the typical expression of anabolic and catabolic genes, and reduced proteoglycan content (Newton Ede and Jones, 2016; Roberts et al., 1993). Analysis of the transcriptional and epigenetics changes in IVD tissue extracted from *Col2-cKO* and littermate control mice demonstrated altered regulation of several anabolic factors and extracellular matrix (ECM) components known to be important for cartilaginous tissues including SOX5/6/9 genes (Liu et al., 2019; Makki et al., 2020). We assayed SOX9, an essential transcriptional regulator of cartilage and of IVD homeostasis in mouse (Henry et al., 2012), which showed strong expression throughout the IVD in littermate control mouse spine at P20 (Fig. 2C-C”). In contrast, *Col2-cKO* mice showed reduced SOX9 expression throughout the IVD, except within the nucleus pulposus (Fig 2D-D”, and G). Collagen type II (COLII) is expressed in the cartilaginous tissues of IVDs with a concentration gradient of high expression in the nucleus pulposus, decreasing outwards the annulus fibrosus, while collagen type I (COLI) is mostly expressed in the annulus fibrosus (Beard et al., 1980; Mader et al., 2016). We observed normal expression of COLII in the IVDs regardless of genotype (Fig. 2E, F), however the typical distribution of COLI positive (COLI+) cells was altered in the *Col2-cKO* mice (Fig. 2H-M). In control IVDs, COLI+ cells are observed in the inner annulus fibrosus, and sparsely in the cartilaginous endplate (red arrows, Fig. 2H-J). However, *Col2-cKO* mutants showed a reduction in COLI+ cells in the inner annulus fibrosus at the convex side of the curvature (Fig. 2K), concomitant with increased COLI+ cells in the inner annulus fibrosus on the concave side (red arrows, Fig. 2L). Interestingly, the growth plate consistently showed increased COLI+ cells in *Col2-cKO* mice (green arrows, Fig. 2M).

Proteoglycans are a broad category of macromolecules characterized by a core protein modified by a diverse array of complex glycosaminoglycan (GAG) chains (Iozzo and Schaefer, 2015) and make up the bulk of ECM of the IVD. We utilized Strong-Anion-Exchange high-pressure liquid chromatography (SAX-HPLC) to quantify proteoglycan species within the IVDs isolated from Cre (-) control and *Col2-cKO* mice at P20. The main GAG species detected are a variety of chondroitin sulfate (CS) species and a minor amount of hyaluronan (HA) species. However, we observed no significant changes in GAG quantity or alterations in specific GAG species (Table S1), demonstrating that loss of *Adgrg6* did not obviously alter the expression of a major class of proteoglycan species in the IVD at the initiation of scoliosis.

In summary, these results show that loss of *Adgrg6* in osteochondral progenitor cells leads to alterations in the typical expression pattern of SOX9+ and COLI+ cells within the IVDs, while alterations of bulk ECM components including COLII and GAG proteoglycans were not greatly affected at the onset of scoliosis. This suggests that even minor alterations in SOX9 expression in the IVD of mice may be sufficient to initiate biomechanical instability leading to disc wedging and driving the onset of spine curvature.

### *Adgrg6* regulates cAMP/CREB signaling dependent gene expression in the IVD

Next, we set out to understand how *Adgrg6* regulates homeostasis of the IVD. Given the established role of *Adgrg6* for control of cAMP signaling in other contexts (Monk et al., 2009; Monk et al., 2011), we first analyzed the expression pattern of Serine-133 phosphorylated CREB (pCREB) --the active form of cAMP response element binding protein (CREB)-- in the spine tissues. pCREB expression is normally enriched throughout cartilaginous elements of the spine, including the cartilaginous endplate and nucleus pulposus (Fig. 3A-A’’ and C). It also expressed in some cells of the periosteum (Fig. 3A’’’ and C). In contrast, *Col2-cKO* mice showed a decrease in pCREB+ cells in these tissues (Fig. 3B-B’’’ and C), suggesting that *Adgrg6* is essential for regulation of pCREB expression in cartilaginous tissues of the spine. We previously demonstrated that *Adgrg6* knockout (KO) ATDC5 chondrogenic cells displayed global disruptions of typical chondrogenic gene expression (Liu et al., 2019). In agreement with our findings in the intervertebral disc we also show that CREB phosphorylation was reduced in *Adgrg6* KO ATDC5 cells compared with unedited control cells (*n=*5 biological repeats) (Fig. 3D, E). Given that *Adgrg6* has been shown to signal through Gs-cAMP in other contexts (Geng et al., 2013; Mogha et al., 2013; Monk et al., 2009), we next asked whether treatment with Forskolin (FSK), a small molecule activator of the adenylyl cyclase, could induce pCREB expression independent of *Adgrg6* function in ATDC5 cells (Fig. 3D, E). We showed that pCREB expression is reduced in *Adgrg6* KO cells compared with in ATDC5 WT cells and that FSK could stimulate pCREB expression in both ATDC5 WT cells and *Adgrg6* KO cells, albeit at much reduced levels after treatment of *Adgrg6* KO cells.

**Figure 3.**
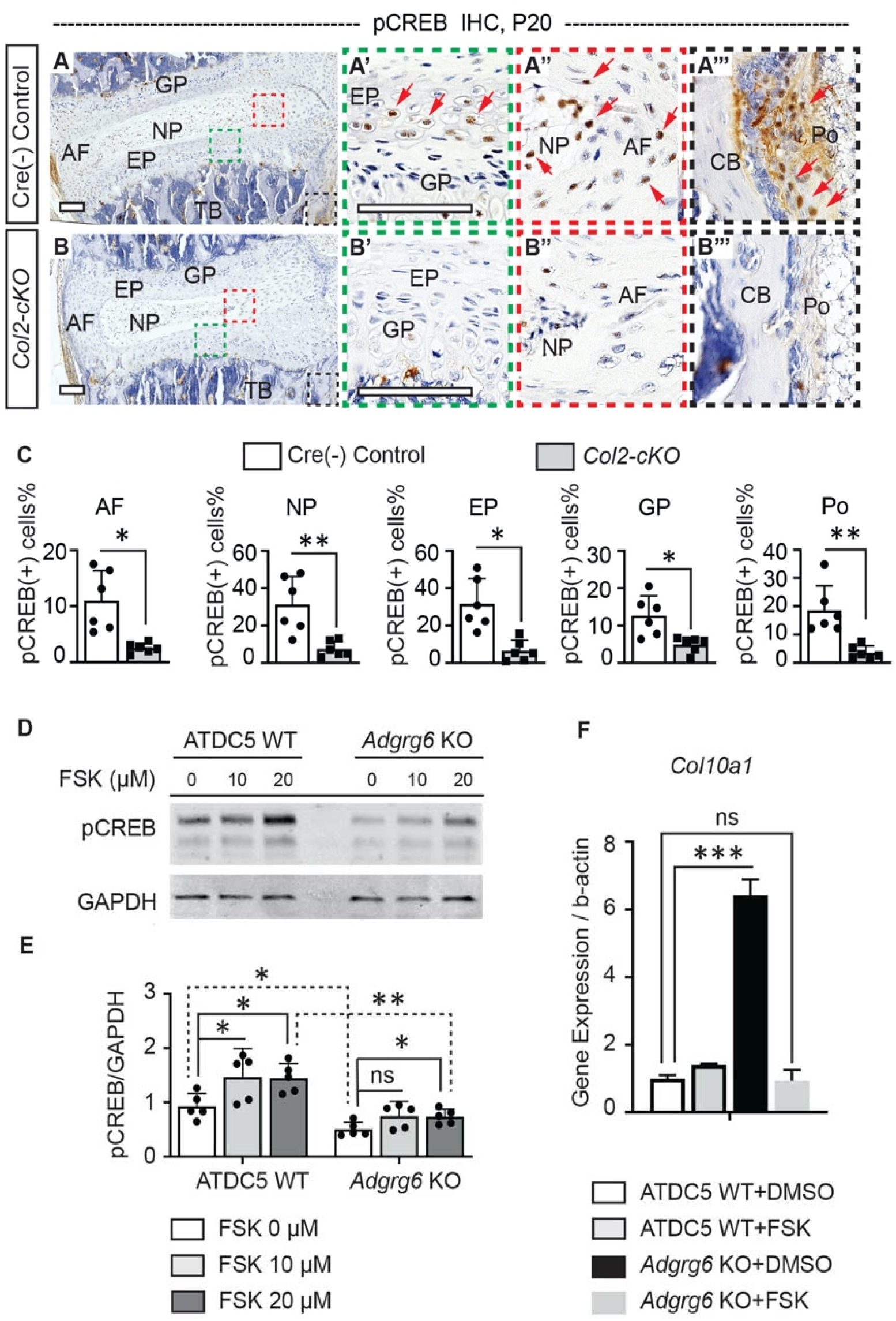
ADGRG6 regulates cAMP/CREB signaling dependent gene expression in cartilaginous lineages. **(A-C)** IHC analyses of pCREB in mouse spine sections at P20. Cre (-) control mice exhibit pCREB positive cell in AF, NP, EP, GP and Po (red arrows, A-A’’’). *Col2-cKO* mutant mice show reduced expression of pCREB in all these tissues (B-B’’’). pCREB positive cells are quantified in (C). *n*=3 mice for each group. Bars were plotted with mean and SD. Each dot represents one IVD analyzed. *: *p*<0.05; **: *p*<0.01, two-tailed Student’s *t* test. ns: not significant. **(D, E)** Representative Western blot image (D) and densitometry of the Western blot images (E) on pCREB in both wild type (WT) and *Adgrg6* KO ATDC5 cell lysates. The expression level of pCREB is significantly decreased in *Adgrg6* KO cells compared with in ATDC5 WT cells. Treatment with 20μM of Forskolin (FSK) for 30min can stimulate pCREB expression in both ATDC5 WT cells and *Adgrg6* KO cells, though the induction level in *Adgrg6* KO cells is significantly lower than that in ATDC5 WT cells (E). *n*=5 biological replicates. Bars are plotted with mean and SD. *: *p*<0.05; **: *p*<0.01, Two-Way ANOVA followed by Tukey test. ns: not significant. **(F)** Real-time RT-PCR analyses of *Col10a1* in ATDC5 WT cells and *Adgrg6* KO cells, cultured with FSK (2μM) or DMSO control for 7 days with maturation medium. The increased expression of *Col10a1* in *Adgrg6* KO cells was rescued by FSK treatment. *n*=3 biological replicates, and the representative result is shown. Bars were plotted with mean and SD. ***: *p*<0.005, One-Way ANOVA followed by Tukey test. ns: not significant. Scale bars: 100μm. *AF- annulus fibrosis, EP- endplate, GP- growth plate, NP- nucleus pulposus, TB-trabecular bone, CB-cortical bone, and Po-periosteum*.

Loss of *Adgrg6* in cartilaginous tissues of the spine led to increased hypertrophy and expression of type X collagen (encoded by *Col10a1*) in cartilaginous endplate and growth plate (Liu et al., 2019). Interestingly, asymmetric expression of type X collagen in vertebral growth plates is observed in AIS patient samples (Wang et al., 2010) and induction of CREB signaling is associated with reduced *Col10a1*expression in primary growth plate chondrocytes (Li et al., 2004). Next, we set out to determine if FSK treatment could rescue altered gene expression outputs observed after loss of *Adgrg6* function. Interestingly, we found that treatment of ATDC5 cells with a low concentration of FSK (2µM) for 7 days was able to restore typical expression of *Col10a1* and stimulate and partially restore *Sox9* expression in *Adgrg6* KO cells (Fig. 3F and S2). While, the dysregulation of other known affected genes, such as decreased expression of *Acan* and *Col2a1* in *Adgrg6* KO cells, was not restored by this approach (Fig. S2). Taken together, these results support a model where *Adgrg6* is necessary for cAMP driven CREB signaling in several cartilaginous tissues, for regulation of a subset of essential genes involved in homeostasis of the IVD.

### ADGRG6 is expressed in the intervertebral disc and supraspinous ligaments

Next, we set out to determine the tissues in which *Adgrg6* function is required for normal spine alignment. To accomplish this, we first assayed the expression pattern of *Adgrg6* in mouse spine using fluorescent *in situ* hybridization, which showed expression in the vertebral growth plate, nucleus pulposus, and the cartilaginous endplate (Fig. 4A-C”). We also observed *Adrgrg6* expression in cells located in the trabecular bone region (Fig. 4A), throughout the annulus fibrosus (Fig. 4C’, C”), and in the connective tissue surrounding the IVD (the outermost annulus fibrosus, white arrows, Fig 4C’), which are recapitulated by immunohistochemistry using a ADGRG6 antibody (Fig. S3A-A”).

**Figure 4.**
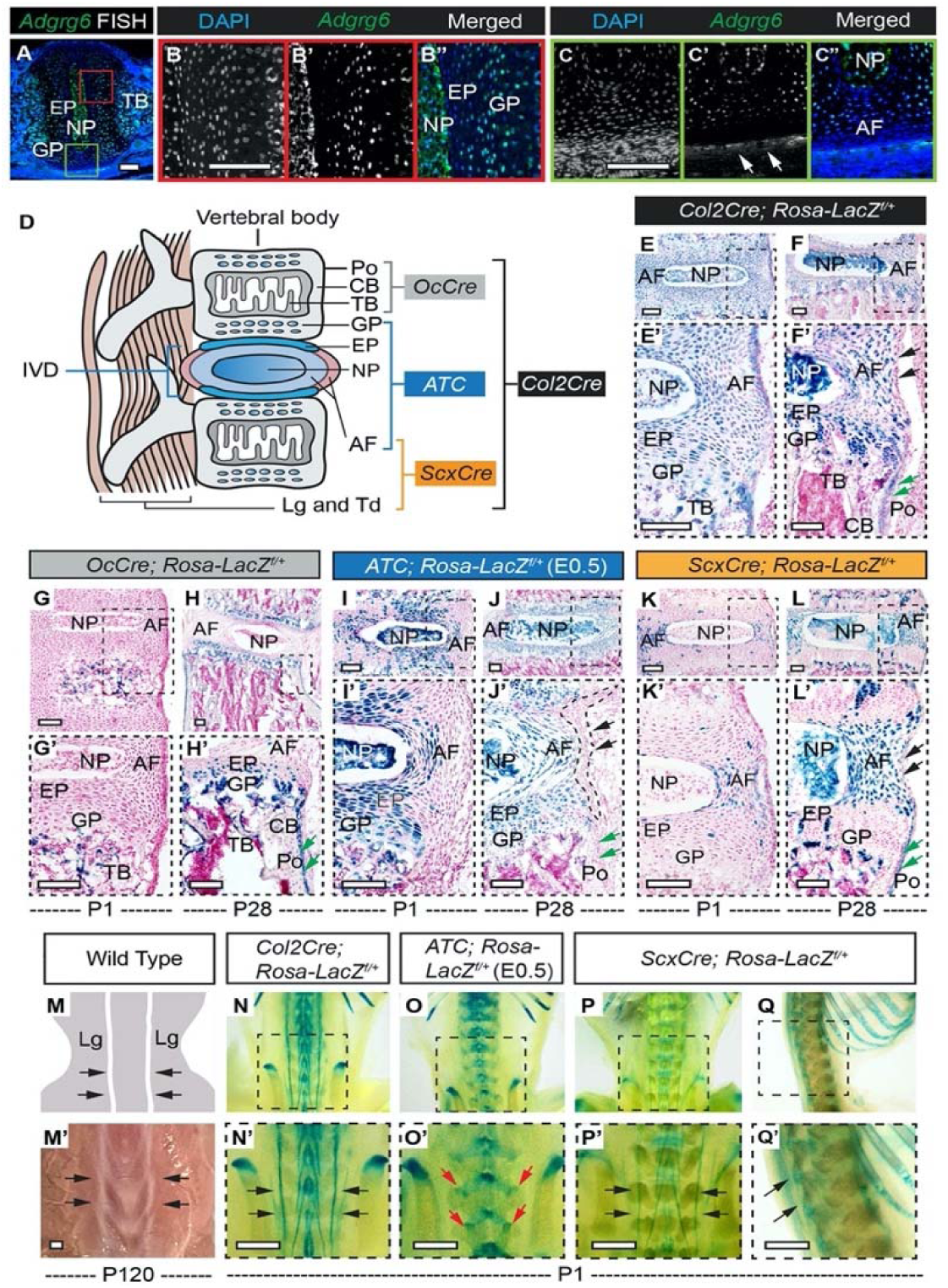
β-galactosidase staining of *Rosa-LacZ* reporter mice recombined with different Cre strains. **(A-C”)** Fluorescent *in situ* hybridization (FISH) analysis of *Adgrg6* on thoracic spine sections of wild type mice at P1. Strong *Adgrg6* signal is detected in the vertebral GP, NP, and in cartilaginous EP (B’, B”). *Adgrg6* is also expressed in TB, AF (A, C’, and C”) and the outmost AF (white arrows, C’). *n*=3 mice for each group. **(D)** An illustration of the spinal tissues targeted by different Cre strains used in this study. Generally, *OcCre* targets bony tissues, *ATC* targets cartilaginous tissues, *ScxCre* targets connective tissues, and *Col2Cre* targets all these tissues. **(E-F’)** β-galactosidase staining of *Col2Cre; Rosa-LacZ^f/+^* spine sections at P1 (E, E’) and P28 (F, F’). *Col2Cre* targets NP, AF, EP and GP, as well as some cells in the bony tissues at both P1 and P28. Note that *Col2Cre* also targets the outmost AF (black arrows, F’) and the periosteum (Po) (green arrows, F’) at P28. **(G-H’)** β-galactosidase staining of *OcCre; Rosa-LacZ^f/+^* spine sections at P1 (G, G’) and P28 (H, H’). *OcCre* targets TB at P1 (G, G’), but targets TB (H’), CB (H’) and Po (green arrows, H’) at P28. *OcCre* also recombines in some hypertrophic cells within the EP and GP (H’). **(I-J’)** β-galactosidase staining of *ATC; Rosa-LacZ^f/+^* (Dox induced from E0.5-P20) spine sections at P1 (I, I’) and P28 (J, J’). *ATC* targets most cells in NP, AF, EP, and GP at both time points. Note that *ATC* did not target the outmost AF (black dash line and arrows, J’), nor the Po (green arrows, J’). **(K-L’)** β-galactosidase staining of *ScxCre; Rosa-LacZ^f/+^* spine sections at P1 (K, K’) and P28 (L, L’). *ScxCre* targets AF at P1 (K’) and P28 (L’), and also recombines in several cells of NP, EP and GP (L’) at P28. Note that *ScxCre* also targets some cells of the outmost AF (black arrows, L’) and the Po (green arrows, L’) at P28. **(M-Q’)** Whole mount β-galactosidase staining of *Rosa-LacZ* reporter mice at P1. An illustration (M) and a bright field image (M’) of the dorsal side of a wild type mouse shows the supraspinous ligaments (black arrows, M, M’). Dorsal view of the whole mount β-galactosidase staining of a *Col2Cre; Rosa-LacZ^f/+^* mouse is shown in (N, N’). Supraspinous ligaments are indicated with black arrows (N’). Dorsal view of the whole mount β-galactosidase staining of an *ATC; Rosa-LacZ^f/+^* mouse (Dox induced from E0.5) is shown in (O, O’). Facet joints are indicated with red arrows (O’). Dorsal and sagittal views of the whole mount β-galactosidase staining of a *ScxCre; Rosa-LacZ^f/+^* mice are shown in (P, P’) and (Q, Q’), respectively. Supraspinous ligaments are indicated with black arrows (P’, Q’). *n*=3 mice in each group. Scale bars: 100μm in A-C and E-L’; 1mm in M’-Q’. *AF- annulus fibrosis, EP- endplate, GPgrowth plate, NP- nucleus pulposus, TB-trabecular bone, CB-cortical bone, Po-periosteum, Lg-ligament, and Td-tendon*.

The *Col2Cre* strain we utilized recombines in osteochondral progenitor cells (Long et al., 2001), which potentially give rise to multiple lineages crucial for building the spine and involved in regulation of spine alignment (Liu et al., 2019; Zheng et al., 2019). To clearly elucidate recombination of the *Col2Cre* strain in the spine we crossed in a *Rosa-LacZ* reporter allele (Soriano, 1999), which showed strong recombination in all the bone- and cartilage-forming lineages of the spine, including the cartilaginous endplate, nucleus pulposus, the entire annulus fibrosus, and the vertebral growth plate (Fig. 4E-F’). This *Col2Cre* strain also targeted the periosteum, some bone-forming cells in the trabecular and cortical bone (Fig. 4E-F’), as well as the ribs and sternum (Fig. S3B, E). Whole mount β-galactosidase staining of a *Col2Cre; Rosa-LacZ^f/+^* mouse at P1 showed that *Col2Cre* also recombined in some connective tissues (*e.g.* ligaments and tendons) along the spine (Fig. 4N, N’), such as the supraspinous ligaments (black arrows, Fig. 4M’, N’). Coronal and transverse sections of the *Col2Cre; Rosa-LacZ^f/+^* mouse spine showed that most cells in the supraspinous ligaments were LacZ positive (Fig. S3H-L), and IHC analyses on adjacent sections showed that these cells also expressed ADGRG6 (Fig. S3I’, J’, and L’). Taken together, these data demonstrate *Adgrg6* is expressed in several elements of the spinal column including cartilaginous, bony and connective tissues which are effectively recombined with *Col2Cre*.

We next sought to genetically dissect which of these tissues require *Adgrg6* function for maintaining spine alignment in mouse. For this, we obtained several CRE driver mouse strains reported to recombine in distinct tissue elements of the spine (Fig. 4D). To assay *Adgrg6* function in bone we used *Osteocalcin-Cre* (*OcCre*) strain to target recombination in mature osteoblasts (Zhang et al., 2002). β-galactosidase staining of spine sections of *OcCre; Rosa-LacZ^f/+^* mice at P1 and P28 showed that *OcCre* specifically targeted mature osteoblasts in the newly forming trabecular bone (Fig. 4G, G’) and at P28, LacZ expression was observed in trabecular bone cortical bone, periosteum, as well as some hypertrophic cells in the vertebral growth plate and cartilaginous endplate (Fig. 4H, H’).

To assay *Adgrg6* function specifically in committed chondrocyte lineages we used a <u>A</u>ggrecan enhancer-driven, <u>T</u>etracycline-inducible <u>C</u>re strain (*ATC*) (Dy et al., 2012; Liu et al., 2019). β-galactosidase staining of skeletal sections from ATC*; Rosa-LacZ^f/+^* mice induced at embryonic (E)0.5 showed robust recombination in cartilaginous tissues of the IVD, including within the cartilaginous endplate, nucleus pulposus, vertebral growth plate, and most cells of the inner annulus fibrosus (Fig. 3I-J’). Interestingly, the outermost annulus fibrosus and the periosteum were not effectively recombined by *ATC* (Fig. 3J’, black arrows and green arrows, respectively). Similar recombination pattern was observed in *ATC; Rosa-LacZ^f/+^* mice when induced at P1 (Liu et al., 2019). Whole mount β-alactosidase staining of *ATC; Rosa-LacZ* mice at P1 showed recombination of *ATC* in the facet joints of the spine (red arrows, Fig. 3O’) and the distal cartilaginous portion of the ribs (red arrow, Fig. S3C), however it did not show robust recombination of the bony portion of the ribs and sternum (yellow arrows, Fig. S3C, F). In summary, both embryonic and perinatal induction of *ATC* driven recombination effectively targets committed cartilaginous tissues, without significant recombination in bone and connective tissues of the spine.

### *Adgrg6* regulates biomechanical properties of dense connective tissues to maintain spine alignment

Dense connective tissues (*e.g.* paraspinal ligaments/tendons) of the spine are effectively targeted by the *Col2Cre* strain (Fig. 4N, N’), but not by the *ATC* strain (Fig. 4O, O’). We hypothesized that these connective tissues may play a synergistic role along with the cartilaginous tissues of the IVD to maintain spine alignment. In addition, the outermost annulus fibrosus (black arrows, Fig. 4F’), as well as some cells in the periosteum (green arrows, Fig. 4F’), which collectively express ADGRG6 (Figure S3A-A”, I’, J’ and L’) are not effectively targeted by the *ATC* strain (Fig. 4J’). To specifically target dense connective tissues of the spine we utilized the *ScxCre* strain which recombines in tendon and ligament progenitor cells (Blitz et al., 2009). Whole mount β-galactosidase staining of *ScxCre;Rosa-LacZ^f/+^* mice at P1 revealed that *ScxCre* targeted the same supraspinous ligaments (black arrows, Fig. 4P, P’), as is observed in the *Col2Cre;Rosa-LacZ^f/+^* mice (black arrows, Fig. 4N, N’). β-galactosidase staining of spine sections from *ScxCre; Rosa-LacZ^f/+^* mice revealed sparse targeting of the annulus fibrosus at P1 (Fig. 4K, K’) but robust targeting of the annulus fibrosus at P28 (Fig. 4L-L’). *ScxCre* also showed sparse recombination of a subset of cells in the nucleus pulposus, endplate, growth plate (Fig. 4L-L’), and the periosteum (green arrows, Fig. 4L’) at P28. Notably, *ScxCre* did not target the bony portion of the ribs or sternum (yellow arrows, Fig. S3D, G), but recombined in some cells of the cartilaginous portion of the ribs (red arrow, Fig. S3D). In summary, we show that *ScxCre* recombines in dense connective tissues of the spine, with only minor contribution to other structural elements of the spine.

### *Adgrg6* is dispensable in osteoblast lineages for the development and alignment of the spine

Abnormal bone quality or osteopenia is frequently associated with AIS (Cheng et al., 2000; Cheng et al., 2001; Lam et al., 2011). To examine whether scoliosis in *Col2-cKO* mice could be further explained by alterations in bone quality we generated *OcCre;Adgrg6^f/f^* (*Oc-cKO*) mice. All *Oc-cKO* mice we assayed showed straight spine alignment at P40 and P120 (*n=*8) (Fig. 5C, E), which was typical of both Cre (-) or Cre (+) littermate controls (Fig. 5A, B, D). Similar results were observed in mice in *Sp7Cre;Adgrg6^f/f^* mice in which *Adgrg6* was ablated in committed osteoblast precursors (Figure S4C, *n*=9). Microcomputed tomography (microCT) analyses revealed normal architecture of the vertebral bodies and comparable bone mass in the thoracic spines of both Cre (-) control and *Oc-cKO* mice at P120 (Fig. 5F-I), which was confirmed by the bone volume analysis (BV/TV) on both control and mutant vertebral bodies (Fig. 5J, *n*=4 for each group). We did not observe obvious histopathological changes of the vertebral bodies or IVDs of the *Oc-cKO* mice at P120 (Fig. S4D, E). Real-time RT-PCR analyses revealed very low expression of *Adgrg6* in the Cre (-) control sample compared with the expression of *Col1a1* (0.2% expression of *Col1a1*), which is the most abundant collagen expressed in bone (Long, 2011) (Fig. 5K). However, the expression of *Adgrg6* was significantly reduced (∼71%) in *Oc-cKO* mice (Fig. 5L). Together these results show that *Adgrg6* has no obvious role in mature osteoblasts during postnatal bone mass accrual or homeostasis and that alterations in bone quality are not driving scoliosis in *Col2-cKO* AIS mouse model.

**Figure 5.**
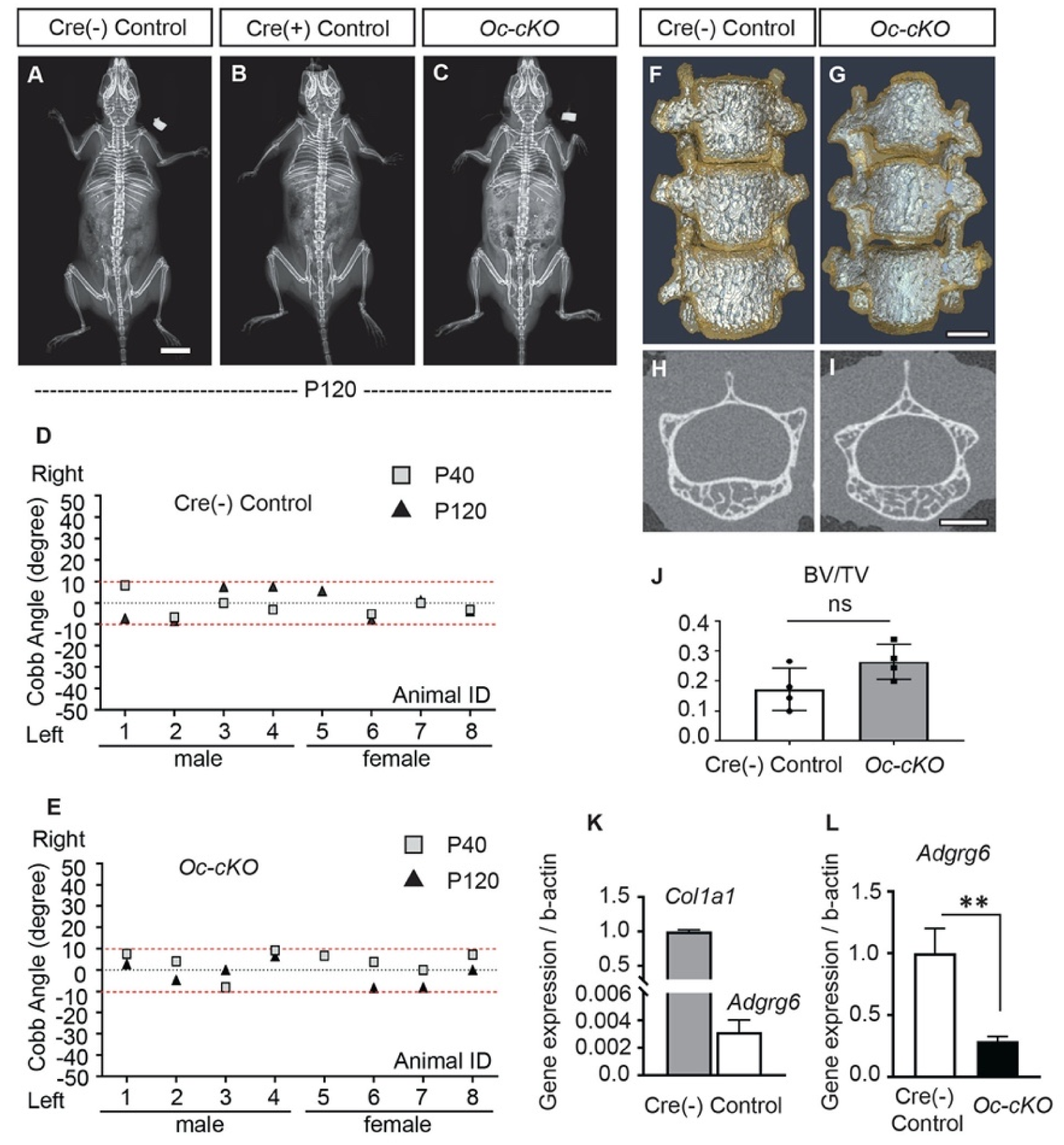
Loss of *Adgrg6* in mature osteoblast lineages is dispensable of AIS development. **(A-C)** Representative X-ray images of Cre (-) control (A), Cre (+) control (B), and *Oc-cKO* mutant (C) mice at P120. **(D, E)** Longitudinal analyses of Cobb angle values of Cre (-) control mice (D) and *Oc-cKO* mice (E) at P40 and P120. For Cre (-) control mice, *n*=8 mice at P40 and P120; for *OcCre; Adgrg6^f/f^* mutant mice, *n*=8 and 7 at P40 and P120, respectively. Thresholds of scoliosis (Cobb angle >10 degree) are indicated with two red dot lines. No *Oc-cKO* mice showed scoliosis at P40 (0/8) or P120 (0/7) (E). **(F-J)** MicroCT scanning of the thoracic region of the spine shows normal morphology of the vertebral bodies in both Cre (-) controls (F) and the *Oc-cKO* mice (G). Transverse sections of the microCT three-dimensional reconstruction of the thoracic vertebral body shows comparable bone mass in the control (H) and mutant mice (I). The bone volume per total volume (BV/TV) of the control and mutant mice was shown in (J). *n*=4 mice for each group. Bars are plotted with mean and SD. ns: not significant. **(K, L)** Real-time RT-PCR analysis of RNA isolated from long bone shows that the expression of *Adgrg6* is very low in bony tissues, compared with the expression of *Col1a1* (K). However, the expression of *Adgrg6* was efficiently knockdown in *Oc-cKO* mice (L). RNA was isolated and pooled from three mice of each experimental group. Bars are plotted with mean and SD. **: *p*<0.01, two-tailed Student’s *t* test. Scale bars: 10mm in A; 1mm in G and I.

### Specific ablation of *Adgrg6* in cartilaginous tissue of the IVD leads to scoliosis

The *ATC* mouse strain allows us to test whether *Adgrg6* has a temporal function in committed cartilaginous in *ATC;Adgrg6^f/f^* (hereafter called *ATC-cKO*) mutant mice, we performed IHC analysis (Fig. S5) and FISH analysis (Liu et al., 2019) on spine sections of both control and *ATC-cKO* mutant mice, and confirmed that ADGRG6/*Adgrg6* expression was effectively knocked-down in cartilaginous tissues of *ATC-cKO* mice using either embryonic or perinatal induction strategies.

Longitudinal X-ray analyses demonstrated that both embryonic and perinatal ablation of *Adgrg6* in committed cartilage tissues of the axial skeleton led to moderate to low penetrance of scoliosis in *ATC-cKO* mice (Fig. 6). Ablation of *Adgrg6* during embryonic development in this way resulted in 25% (*n=*16) and 16.7% (*n=*12) of *ATC-cKO* mice showing scoliosis at P20 and P180 respectively (Fig. 6A-C and H). Cobb angle measurements revealed curve severity between 11° to 43° in scoliotic mutant mice (Fig. 6H). Ablation of *Adgrg6* during perinatal development in *ATC-cKO* mice showed no evidence of scoliosis when assayed at P20 (*n=*8), while a single *ATC-cKO* mutant mouse (*n=*8) showed late-onset scoliosis when assayed at P180 with a curve of 23.8° (Fig. 6 D-F and J). None of the Cre (-) littermate control mice showed evidence of spine deformity at either P20 or P180 (Fig. 6G, I). Histological analyses of the *ATC-cKO* mutant spine were performed previously, showing altered extracellular matrix component expression and endplate-oriented disc herniations during later stages of adult development (8 months)(Liu et al., 2019). Altogether this suggested that loss of *Adgrg6* in committed cartilages of the axial skeletal is sufficient to generate scoliosis, although with a much lower penetrance compared with removal of *Adgrg6* in whole spine (*Col2-cKO*)(Fig. 1L). These results suggest that the pathogenesis of this model of postnatal scoliosis is affecting processes of spine biology established during embryonic development.

**Figure 6.**
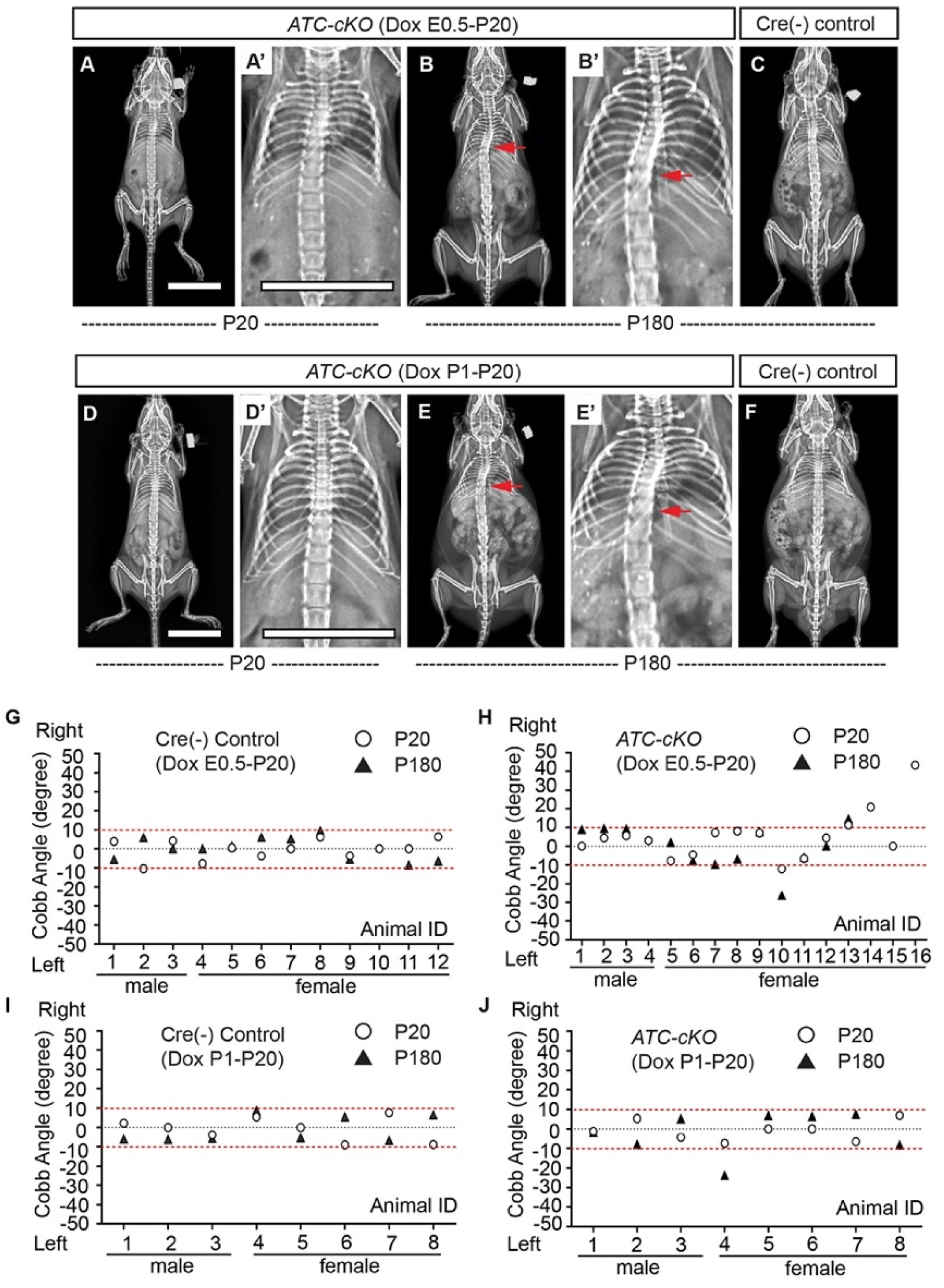
Ablation of *Adgrg6* in cartilaginous tissues leads to scoliosis in mouse. **(A-F)** Representative X-ray images of Cre (-) control and *ATC-cKO* mutant mice at P20 and P180. *ATC-cKO* mice (Dox induction from E0.5-P20) analyzed at P20 and P180 are shown in (A, A’) and (B, B’), respectively. *ATC-cKO* mice (Dox induction from P1-P20) analyzed at P20 and P180 are shown in (D, D’) and (E, E’), respectively. Corresponding Cre (-) control mice analyzed at P180 were shown in (C) and (F). Scoliosis is indicated with red arrows in (B, B’) and (E, E’). **(G-J)** Longitudinal analyses of Cobb angle values of Cre (-) control mice and *ATC-cKO* mice at P20 and P180. For embryonic induction (E0.5-P20), *n*=12 mice for Cre (-) controls at P20 and P120; *n*=16 and 12 for *ATC-cKO* mice at P20 and P180, respectively. For perinatal induction (P1-P20), *n*=8 mice for Cre (-) controls at P20 and P180; *n*=8 for *ATC-cKO* mice at P20 and P180. Thresholds of scoliosis (Cobb angle >10 degree) are indicated with two red dot lines. Scale bars: 10mm.

To test whether this affect was due to a real biological effect or due to an idiosyncratic property of the *ATC* strain, we utilized an alternative *Col2CreER^T2^* strain, to target cartilaginous tissues of the spine postnatally (Chen et al., 2007). Next, we performed two temporal induction Tamoxifen strategies: (i) perinatal induction from P1-P5; and (ii) postnatal induction from P14-P18. β-galactosidase staining of spine sections of *Col2CreER^T2^; Rosa-LacZ^f/+^* mice at P28 showed that perinatal induction (P1-P5) led to near complete recombination in cells of cartilaginous endplate, vertebral growth plate, and most cells of the inner annulus fibrosus but not the outer annulus fibrosus (Fig. S6A, A’), similar to the recombination profile of *ATC* (Fig. 4I-J’), except that *Col2CreER^T2^* did not target the nucleus pulposus (Fig. S6A). Unfortunately, postnatal induction (P14-P18) in *Col2CreER^T2^;Rosa-LacZ^f/+^* mice only led to partial recombination within endplate and growth plate, and some cells of the inner annulus fibrosus, while the outer annulus fibrosus and nucleus pulposus were not targeted (Fig. S6B, B’). We found no evidence of scoliosis in *Col2CreER^T2^; Adgrg6^f/f^* (hereafter called *Col2TM-cKO*) mutant mice after perinatal induction (P1-P5) at P30 0.0% (*n=*8) (Fig. S6 C, C’ and J). However, a single (*n=*7) *Col2TM-cKO* mice presented with very mild late-onset scoliosis (12.4°) at P90 (Fig S6D, D’ and J). The lower penetrance of scoliosis in *Col2TM-cKO* mice was comparable with the results we observed in the *ATC-cKO* mice induced from P1-P20 (Fig. 6J). We also found no evidence of scoliosis in *Col2TM-cKO* mice induced at P14-P18 at either P30 or P90 (*n=*5 for *Col2TM-cKO* and *n=*4 for controls, respectively) (Fig S6F-H, K, and L). Taken together, our results confirmed that *Adgrg6* plays only a minor role in postnatal regulation of cartilaginous tissues for maintaining spine alignment in mouse. However, our results showed a slight increase in the incident of scoliosis in the *ATC-cKO* mice with embryonic deletion of *Adgrg6*, compared to that with perinatal deletion. This further supports a model where the contribution of cartilaginous elements of the spine for perinatal spine alignment are chiefly established during embryonic development in mouse.

### Ablation of *Adgrg6* in dense connective tissues leads to late-onset scoliosis

Connective tissue disorders such as Ehlers-Danlos and Marfan syndromes display scoliosis as a common symptom (Hoffjan, 2012) and generalized joint hypermobility is a risk factor for developing AIS (Haller et al., 2018). To address a specific role of *Adgrg6* in dense connective tissues of the spine we generated *ScxCre;Adgrg6^f/f^* (*Scx-cKO*) mutant mice, which were indistinguishable from littermate controls at birth. Longitudinal X-ray analyses revealed that 55.6% (*n=9*) of *Scx-cKO* mice exhibited scoliosis at P120, with curve severity ranged between 12° to 30° (Fig. 7B, B’, and F). Interestingly, none of these mice showed scoliosis at P40 (*n=*9) (Fig. 7A, A’, and F). No scoliosis was observed in Cre (-) or Cre (+) littermate control mice at either P40 or P120 (Fig. 7C-E). Histological analysis revealed some mildly wedged IVDs and shifted nucleus pulposus within the scoliotic curve of the *Scx-cKO* mutant spine at P120 (Fig. S7C), with IVDs outside of the curve region displaying typical morphology (Fig. S7A, B). Altogether this demonstrates that dense connective tissues of the spine are essential for maintaining spine alignment later during adult development (after P40) and may act synergistically with the IVD to promote typical spine alignment during postnatal development in mouse.

**Figure 7.**
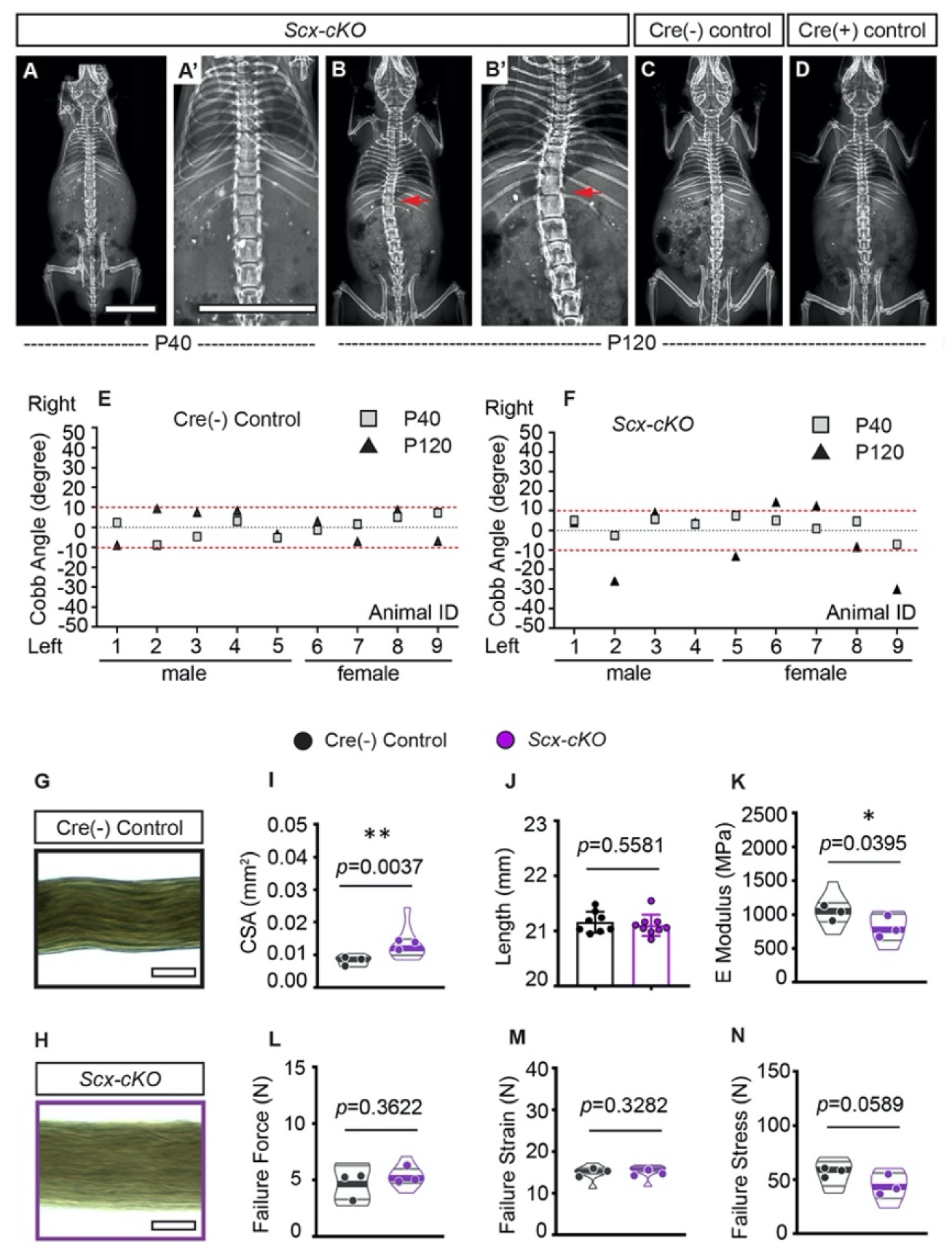
Ablation of *Adgrg6* in dense connective tissues leads to late-onset scoliosis and compromised biomechanical properties of the tendons. **(A-D)** Representative X-ray images of Cre (-) control and *Scx-cKO* mutant mice. *Scx-cKO* mice analyzed at P40 and P120 are shown in (A, A’) and (B, B’), respectively. Cre (-) control and Cre (+) control mice analyzed at P120 are shown in (C) and (D), respectively. Scoliosis is indicated with red arrows in (B, B’). **(E, F)** Longitudinal analyses of Cobb angle values of Cre (-) control mice and *Scx-cKO* mice at P40 and P120. For Cre (-) control mice, *n*=9 mice at both P40 and P120; for *ScxCre; Adgrg6^f/f^* mutant mice, *n*= 9 at both P40 and P120. Thresholds of scoliosis (Cobb angle >10 degree) are indicated with two red dot lines. **(G-N)** Biomechanical characterization of Cre (-) control and *ScxcKO* mutant tendons at 12 weeks. Representative phase-contrast images of tail fascicles isolated from Cre (-) control and *Scx-cKO* mutant mice are shown in (G) and (H). Quantification of fascicles’ cross-sectional area (CSA) (I), initial length (J), elastic modulus (E Modulus) (K), failure force (L), failure strain (M), and failure stress (N) are also shown. *n*= 8-9 fascicles isolated from 3 mice. For (I) and (K-N), *n*= 8-9 fascicles (violin plots) isolated from 3 different mice (close circles). For J, *n*=8 and 9 fascicles isolated from Cre (-) control mice or *Scx-cKO* mutant mice, respectively. Bars are plotted with mean and SD. Cross-sectional area (I) and elastic modulus (K) were significantly different between control and mutant groups. *: *p*<0.05, **: *p*<0.01, unpaired *t* test with Welch’s correction for (J, K, L and N), and with Mann-Whitney test for non-normally distributed data (I and M). Scale bars: 10mm in A, A’; 100μm in G, H.

### Ablation of *Adrgrg6* compromises the biomechanical properties of tendons

Next, to determine if the loss of *Adgrg6* alters the mechanical properties we used tendon fascicles isolated from tails of 12-week-old *Scx-cKO* mice and Cre(-) controls as a proxy for dense connective tissues controlling spine stability. Tail tendons originate from the same *Scx*- expressing somatic progenitor population in the syndetome as axial tendons and ligaments (Murchison et al., 2007). Moreover, the fascicle represents the tissue-level “functional unit” of tendons, and can easily be extracted as an individual, intact unit for highly-reproducible characterization of structure-function relationships (Wunderli et al., 2020). We examined each fascicle to measure the external diameter, and the cross-sectional area was calculated based on these measurements (Fig. 7G, H). We found that the total cross-sectional area was significantly increased in *Scx-cKO* tendons compared to Cre (-) controls (*P*=0.0037) (Fig. 7G, H and I). Micromechanical tensile testing revealed that deletion of *Adgrg6* resulted in mechanically weaker tendons with significantly lower elastic moduli (*P*=0.0395), and trend of reduced failure stresses (*P*=0.0589) (Fig. 7K, N). We observed no differences in failure force or failure strain (Fig. 7L, M). This data suggests that ablation of *Adgrg6* in *Scx*-expressing cells compromises the biomechanical properties of dense connective tissues, which may potentially contribute to the progression of the late-onset scoliosis in *Scx-cKO* mice. Altogether this study demonstrates the essential role of *Adgrg6* and its tissue-level function within the cartilaginous tissues and dense connective tissue of the spine and highlights its role for the maintenance of homeostatic signaling and biomechanical properties of these spine elements to regulate postnatal spine alignment in mouse.

## Discussion

AIS is the most common pediatric disorder worldwide, yet the underlying causes remain a persistent challenge for pediatric medicine. A wide range of hypotheses have been proposed to explain its origins, including deformities in structural skeletal elements, neurological abnormalities, and altered biomechanical loading of the spine (Cheng et al., 2015; Latalski et al., 2017; Newton Ede and Jones, 2016). While, these is a wide range of animal models proposed for understanding the pathogenesis of AIS, with differing levels of validity (Liu and Gray, 2018), we suggest that the susceptibility of scoliosis observed in *Adgrg6* conditional mutant mice provide unique models of AIS with construct validity (Willner, 1984) by modeling both the natural history of this spine disorder (this work) after disruption of a gene associated with AIS in humans (Kou et al., 2013; Kou et al., 2018). Next, we demonstrate that alterations in *Adgrg6* signaling specifically in cartilaginous and dense connective tissues, but not in the bone-forming lineages, are key drivers of scoliosis. Further, our results suggest that stimulation of cAMP signaling may provide pathways for therapeutic interventions of AIS.

There is good agreement of evidence supporting the involvement of cartilaginous tissues in the etiology of scoliosis. For instance, patients with AIS frequently exhibit wedging of IVDs associated with shifted nucleus pulposus to the convex side of the curve (Little et al., 2016), which is precisely recapitulated in *Col2-cKO* mutant mice. T2-weighted magnetic resonance imaging signal distributions in the IVDs can discriminate severity of AIS (Gervais et al., 2012), and the vertebral growth plate abnormities are frequently found to locate near the apex of the curve (Day et al., 2008), suggesting that alterations of cartilaginous tissues may proceed the initiation of scoliosis. In agreement, AIS patients exhibit IVD degeneration including disorganization of chondrocytes in the vertebral growth plate (Day et al., 2008), reduced proteoglycan contents in the endplate and nucleus pulposus (Shu and Melrose, 2018; Urban et al., 2001), and altered distribution of collagen fibers in the annulus fibrosus (Akhtar et al., 2005). Some patients also exhibit increased incidence of Schmorl’s nodes (endplate-oriented disc herniations) (Buttermann and Mullin, 2008). In agreement, these phenotypes are observed in both *Col2-cKO* and *ATC-cKO* mice mutant mice reported here and in our previously (Liu et al., 2019). For example, at the initiation of scoliosis (P20-P40) we observed altered gene expression, disorganized collagen distribution, and signs of disc hypertrophy and disc degeneration in the spines of young *Col2-cKO* or *ATC-cKO* mice (Liu et al., 2019). During the progression of scoliosis (P120-8 months), we observed disorganization of endplate and growth plate cells, as well as disc degeneration and endplate-oriented disc herniation (Liu et al., 2019). All these observations support a model that cartilaginous tissues of the IVD play a crucial role during both initiation and progression of scoliosis.

An interesting question remains as to how *Adgrg6* regulates homeostasis of cartilaginous tissues. We previously showed that *Adgrg6* maintains IVD homeostasis in part via negative regulation of STAT3 signaling (Liu et al., 2019). Here, we demonstrate that *Adgrg6* is also essential for cAMP/CREB signaling. Importantly, cAMP signaling has been shown to have chondroprotective effects in cartilage, stimulating expression of several anabolic markers, such as *Col2a1* and *Acan* (Kosher et al., 1986; Malemud et al., 1986), and blocking matrix metalloproteinase-mediated cartilage degeneration in articular cartilage (Karsdal et al., 2007). Moreover, cAMP signaling is known to positively regulate SOX9, which is a master regulator for both development and maintenance of cartilaginous tissues of the spine (Akiyama et al., 2002; Henry et al., 2012). Specifically, cAMP signaling stimulates phosphorylation of SOX9 protein, which enhances its DNA binding affinity and transcriptional activity (Huang et al., 2001; Huang et al., 2000). Moreover, CREB has also been shown to bind to *SOX9* promoter and induce its expression (Kanazawa et al., 2014; Piera-Velazquez et al., 2007). In agreement, we show that loss of *Adgrg6* function regulates the expression pCREB and SOX9 expression in the IVD and periosteum, partially via stimulation of cAMP. Further, identification of downstream effectors of cAMP/CREB signaling in structural elements of the spine may uncover new susceptibility loci involved in AIS.

The onset of idiopathic scoliosis usually occurs during adolescence in humans (Cheng et al., 2015). *Adgrg6* appears to be largely dispensable for processes crucial for overall spine patterning and formation of the IVDs (Liu et al., 2019). Here, we showed that ablation of *Adgrg6* in the IVD during embryonic development increased the penetrance of scoliosis from less than 15% to about 25%. Therefore, our observations suggest that embryonic deletion of *Adgrg6* leads to subtle structural and/or biomechanical changes of these structural elements of the spine, which in turn may lead to increased susceptibility of scoliosis during periods of rapid skeletal growth.

In contrast to a recent report (Sun et al., 2020), we found that *Adgrg6* has no obvious functional role in bone forming osteoblasts for skeletal morphology. We observed no spinal malformations nor apparent limb or growth phenotype in either *Oc-cKO* mice or *Sp7-cKO* mice (Fig. 5A-C and Fig. S4A-C). Sun *et al*. also reported that scoliosis was not present in *OsxCre;Gpr126^f/f^* mice (equivalent to our *Sp7-cKO* mice) at P120 (Sun et al., 2020). However, in contrast with our findings, Sun *et al*. reported that deletion of *Adgrg6* (*Gpr126*) with *OsxCre*, but not *LysmCre* (which targets osteoclast) or *Col2Cre*, resulted in a significant reduction in overall body size and the bone length. The authors attributed this effect primarily due to delayed osteoblast differentiation (Sun et al., 2020). We and others have shown that *Col2Cre* recombines in early osteochondral progenitor cells that give rise to both osteoblasts and chondrocytes (Liu et al., 2019; Long et al., 2001). On the other hand, *OsxCre* (*Sp7Cre*) is activated in many cells/tissues in addition to osteoblasts, including but not limited to hypertrophic chondrocytes, adipocytes, vascular smooth muscle, perineural and stromal cells in the bone marrow, olfactory glomerular cell, and intestinal epithelial cells (Chen et al., 2014; Liu et al., 2013). Therefore, the presence of a phenotype only in the *OsxCre* but not *Col2Cre* mutant mice could be due to an unidentified cell type or is the effect of the *OsxCre* transgene itself (Davey et al., 2012; Huang and Olsen, 2015; Wang et al., 2015). In summary, while we provide evidence against a role of *Adgrg6* in osteoblasts, we suggest that additional studies are warranted to help allay these divergent results.

The penetrance of scoliosis at P40 in the *Scx-cKO* mice, targetd only dense conn3ective tissues of the spine, is comparable with the penetrance of early-onset scoliosis (before P20) in *Col2-cKO* mice, which targeted all tissues of the spine. Connective tissues, including ligamentous structures and tendon insertions of the spine, along with paraspinal muscles play a key role in controlling spinal stability (Bogduk, 2016). Not surprisingly, defects in connective tissues are reported in a range of scoliosis conditions. For instance, reduced fiber density and non-uniform distribution of elastic fibers were observed in the ligamentous tissues in some AIS patients compared with the healthy individuals (Hadley-Miller et al., 1994). Moreover, connective tissue disorders such as Marfan’s syndrome also commonly display scoliosis (Glard et al., 2008) and rare variants in the Marfan’s associated fibrillin genes *FBN1/2* are have been associated with AIS in humans (Buchan et al., 2014; Sheng et al., 2019). We show that *Scx-cKO* mice exhibit reduced elastic moduli of the tendon fascicles of the tail, which suggests that scoliosis in these *Adgrg6* mutant mouse models of AIS may be largely driven by biomechanical instability of the paraspinal tendons and ligaments. Altogether these results suggest that regulation of biomechanical properties of dense connective tissues may underscore a common pathogenesis of scoliosis observed in between AIS and other connective tissue disorders.

Altogether, our study provides evidence that cartilaginous and connective tissues may serve as origins for some forms of scoliosis, and support the regulatory role and therapeutic value of *Adgrg6*/cAMP signaling for future treatment approaches aimed at halting the initiation and progression of AIS in humans.

## Materials and Methods

### Mouse strains

All animal studies were conducted according to institutional and national animal welfare guidelines, and were approved by the Institutional Animal Care and Use Committee at The University of Texas at Austin (protocol AUP-2018-00276). All mouse strains were described previously, including *Adgrg6^f/f^* (Taconic #TF0269) (Mogha et al., 2013), *Rosa-LacZ* (Soriano, 1999); *ATC* (Dy et al., 2012), *Col2Cre* (Long et al., 2001), *Col2CreER^T2^* (Chen et al., 2007); *ScxCre* (Blitz et al., 2009), *OcCre* (Zhang et al., 2002), and *Sp7Cre* (Rodda and McMahon, 2006). Doxycycline (Dox) was administered to *ATC-cKO* and littermate controls by two strategies: (i) inducing from embryonic day (E) 0.5 to postnatal day (P) 20 by ad libitum feeding of Dox-chow (Test Diet, 1814469) to plugged isolated females, and supplemented via intraperitoneal (IP) injections of the pregnant dames with Dox (Sigma D9891) once/week (10mg/kg body weight) throughout the pregnancy until the pups were weaned at P20; (ii) inducing from P1-P20 by ad libitum feeding of Dox-chow to the mothers after the pups were born, and supplemented with IP injections of the mothers with Dox once/week (10mg/kg body weight) until the pups were weaned at P20. Tamoxifen (Sigma, T5648) was dissolved in filter-sterilized corn oil at a concentration of 20mg/ml, and administered to *Col2TM-cKO* and Cre(-) littermate controls by two strategies: (i) inducing from P1-P5 via injection into the milk dot of the pups once/day for five consecutive days (20µl at P1, 30µl at P2 and P3, and 40µl at P4 and P5); (ii) inducing from P14-P18 via IP injections once/day (1mg/10g) for five consecutive days. *ATC; Rosa-LacZ^f/+^*, *Col2CreER^T2^; Rosa-LacZ^f/+^*, and the corresponding Cre(-) littermate controls were induced with the same strategies.

### Analyses of mice

Radiographs of mouse skeleton were generated using a Kubtec DIGIMUS X-ray system (Kubtec T0081B) with auto exposure under 25 kV. Cobb angle was measured on high-resolution X-ray images with the software Surgimap (https://www.surgimap.com), as previously described (Cobb, 1958).

Histological analysis was performed on thoracic (T5-T12) spines fixed in 10% neutral-buffered formalin for 3 days at room temperature, followed by 5 days of decalcification in Formic Acid Bone Decalcifier (Immunocal, StatLab). After decalcification, bones were embedded in paraffin and sectioned at 5 μm thickness. Alcian Blue Hematoxylin/Orange G (ABH/OG) and Safranin O/Fast Green (SO/FG) staining was performed following standard protocols (Center for Musculoskeletal Research, University of Rochester). Immunohistochemical analyses were performed on paraffin sections with traditional antigen retrieval and colorimetric developed development methodologies with the following primary antibodies: anti-SOX9 (Chemicon AB5535, 1:200), anti-Collagen II (Thermo Scientific MS235B, 1:100), anti-GPCR GPR126 (ADGRG6) (Abcam, ab117092, 1:100), anti-Collagen I (Abcam, ab138492, 1:1000), and anti-phospho-CREB (Ser133) (Cell Signaling 87G3, #9198, 1:200). The β-galactosidase staining was performed on frozen sections as previously described with modifications (Liu et al., 2015). Briefly, mouse spines or P1 pups were harvested and fixed in 10% neutral-buffered formalin for 2 hours or overnight at 4°C, respectively, and thoroughly washed in 1xPBS. Spines were decalcified with 14% EDTA at 4°C for 1 week or overnight, washed in sucrose gradient, embedded with Tissue-Tek OCT medium, snap-frozen in liquid nitrogen, and sectioned at 10μm with a Thermo Scientific HM 550 cryostat. Both spine sections and P1 pups were stained with X-gal (0.5mg/ml) at 37°C overnight protected from light. P1 pups where cleared in 50% glycerol (in 1xPBS and 1% KOH) for 2 weeks at room temperature with gentle rocking, and stored in 80% glycerol.

MicroCT analysis was performed on thoracic regions of the spine (T5-T12) in control and *Oc-cKO* mutant mice at P120. The spines were fixed in 10% neutral-buffered formalin for 3 days at room temperature, thoroughly washed, and scanned on an Xradia at 100 kV at 9.65 μm true voxel resolution at the University of Texas High-Resolution X-ray Computed Tomography Facility (http://www.ctlab.geo.utexas.edu).

### GAG analysis

8-10 pieces of intervertebral disc and adjacent vertebral growth plates were isolate from three Cre(-) control mice and two *Col2-cKO* mutant mice at P20, and snap frozen in liquid nitrogen. The samples were then ground by pestle in Pronase buffer (0.1M Tris-HCl, pH 8.0, containing 2mM CaCl_2_ and 1% Triton X-100), homogenized by sonication (Barnstead probe homogenizer) for 30 sec, and suspended in 2.5ml of Pronase buffer. Samples were digested with Pronase (0.8mg/ml) for 24h at 50 °C with gentle shaking. After 24h, a second aliquot of 1.6mg Pronase was added, and the samples were digested for another 24h, followed by incubation at 100 °C for 15min to inactivate the reaction. The samples were then adjusted with 2mM MgCl_2_ and 100mU benzonase, incubated at 37 °C for 2h, and centrifuged at 4000g for 1h. The supernatant was applied to a DEAE-Sepharose-micro column, washed with loading buffer (0.1M Tris-HCl, pH 8.0) and washing buffer (pH Acetate buffer), and then eluted in 2M ammonium acetate. The acetate salt was removed via lyophilization and the sample was reconstituted in deionized water.

These isolated GAG materials were further digested with either Chondroitinase ABC or Hyaluronidase from *Streptomyces Hyalurolyticus* at 37°C for over 24h, followed by incubation at 100°C for 5 minutes to inactivate the enzymes. Samples were centrifuged at 14,000 rpm for 30 minutes before introduction to the HPLC.

SAX-HPLC was carried out on an Agilent system using a 4.6×250 mm Waters Spherisorb analytical column with 5 μm particle size at 25°C. A gradient from low to high salt was used to elute the GAG disaccharides. The flow rate was 1.0 mL/min and the injection of each sample was 10 μl for the positive controls and samples. Detection was performed by post-column derivatization, and fluorescence detection. Commercial standard disaccharides (Dextra Laboratories) were used for identification of each disaccharide based on elution time, as well as calibration. A positive control digestion of Hyaluronan was used for Hyaluronidase digestions.

### Mechanical testing

Three 12-week-old Cre(-) control mice and *Scx-cKO* mutant mice were euthanized and stored in a -20C° freezer until needed. On the day of mechanical testing, tails were thawed and dissected from the body using surgical blades. Fascicles were extracted by holding the tail from its posterior end with surgical clamps, and gently pulling off the skin until bundles of fascicles were exposed. Each fascicle was examined under a microscope (Motic AE2000, 20x magnification) for visible signs of damage, and for measuring the external diameter. Specimens with frayed ends, visible kinks or diameters less than 90µm were discarded. Micromechanical tensile testing was performed using a custom-made horizontal uniaxial test device to generate load-displacement and load-to-failure curves (10N load cell, Lorenz Messtechnik GmbH, Germany).

Briefly, two-centimeter fascicle specimens were carefully mounted and kept hydrated in PBS, as previously described (Fessel and Snedeker, 2011). Each fascicle underwent the following protocol: pre-loading to 0.015 N (L0: 0% strain), 5 cycles of precondition to 1% L0, an additional 1% strain cycle to calculate the tangential elastic modulus, and then ramped to failure to 20% strain under a predetermined displacement rate of 1L were processed using a custom-written MATLAB script (Matlab® R2018b, v.9.5.0.944444, MathWorks, Inc.). Tangent elastic moduli were calculated from the linear region of stress-strain curves (0.5-1%). Nominal stress was estimated based on the initial cross-sectional area. Cross-sectional area was calculated from microscopic images assuming fascicles have perfect cylindrical shape.

### Cell culture

The *Adgrg6* KO ATDC5 cell line was generated in our lab as previously described (Liu et al., 2019). Both wild type ATDC5 cells (Sigma, 99072806) and *Adgrg6* KO ATDC5 cells were maintained in DMEM/F-12 (1:1) medium (Gibco, 11330032) supplemented with 5% FBS and 1% penicillin/streptomycin. Cells were cultured in 24-well cell culture plate and maturated in DMEM/F-12 (1:1) medium supplemented with 5% FBS, 1% penicillin/streptomycin, 1% ITS premix (Corning, 354352), 50 μg/ml ascorbic acid, 10nM dexamethasone, and 10ng/ml TGF-β3 (Sigma, SRP6552) for 7 days, with stimulation of 2uM Forskolin (FSK, Sigma F6886) or DMSO control. After 7 days, cells were proceeded with RNA isolation and real-time RT-PCR.

### RNA isolation and real-time RT-PCR

To confirm the reduction of *Adgrg6* expression in osteoblast lineages, we isolated total RNA from fresh frozen, pulverized long bones at P120. Total RNA was isolated using a RNeasy mini kit (Qiagen 74104). Reverse transcription was performed using 500 ng total RNA with an iScript cDNA synthesis kit (Bio-Rad). Real-time RT-PCR (qPCR) analyses were performed as previously described (Liu et al., 2015). Gene expression was normalized to β-actin mRNA and relative expression was calculated using the 2-(ΔΔCt) method. The following primers were used to check gene expression: β-actin: AGATGTGGATCAGCAAGCAG/GCGCAAGTTAGGTTTTGTCA; *Col1a1*: GCATGGCCAAGAAGACATCC/CCTCGGGTTTCCACGTCTC; *Adgrg6*: CCAAAGTTGGCAATGAAGGT/GCTGGATCAGGTAGGAACCA; *Col10a1*: CTTTGTGTGCCTTTCAATCG/GTGAGGTACAGCCTACCAGTTTT; *Col2a1*: ACTGGTAAGTGGGGCAAGAC/CCACACCAAATTCCTGTTCA; and *Acan*: CGTGTTTCCAAGGAAAAGGA/TGTGCTGATCAAAGTCCAG. Real-time RT-PCR efficiency was optimized and melting curve analyses of products were performed to ensure reaction specificity.

### Western blotting

Both wild type and *Adgrg6* KO ATDC5 cells were treated with 10µM or 20µM Forskolin for 30min before protein extraction. Total proteins were extracted from cells with the M-PER Mammalian Protein Extraction Reagent (Sigma, GE28-9412-79) supplemented with protease and phosphatase inhibitors (Roche). 10mg of protein from each sample was resolved by 4-15% SDS-polyacrylamide gel electrophoresis and transferred to the nitrocellulose membranes. Western blots were then blocked with LI-COR blocking buffer and incubated overnight with primary antibodies anti-pCREB (Ser133)(Cell Signaling 87G3, #9198, 1:1000) and anti-GAPDH (Cell Signaling, #2118, 1:2500) at 4°C with gentle rocking. The next day western blots were detected with the LI-COR Odyssey infrared imaging system.

### Statistics

Statistical analyses were performed in GraphPad Prism 8.4.3 (GraphPad Software Inc, San Diego, CA). For Cobb angle and gene expression analysis to compare two or more experimental groups, 2-tailed Student’s *t*-test, one-way ANOVA, or two-way ANOVA followed by Turkey HSD test were applied as appropriate. For mechanical testing, normality of distribution was checked using Shapiro-Wilk test. Unpaired Student’s *t*-test with Welch’s correction or Mann-Whitney U-test were used for two-group comparison of parametric and non-parametric data, respectively. A *p* value of less than 0.05 is considered statistically significant.

## Acknowledgements

We thank John Wallingford and Matthew Hilton for comments prior to publication. We thank Drs. Fanxin Long for sharing *Col2Cre* mice, Dr. Véronique Lefebvre for sharing *ATC* mice, Dr. Matthew Hilton for sharing *Col2CreER^T2^* mice, and Dr. Ronen Schweitzer for sharing *ScxCre* mice. We thank Dr. Parastoo Azadi and Ms. Stephanie Archer-Hartmann at the Complex Carbohydrate Research Center for performing the GAG analysis. This study was supported by the National Institute of Arthritis and Musculoskeletal and Skin Diseases of the National Institutes of Health under Award Number R01AR072009-01 (R.S.G.), R01AR071967 and R01AR076325 (C.M.K), and F32AR073648 (Z.L.). It was also supported in part by the National Institutes of Health-funded Research Resource for Integrated Glycotechnology (NIH Award Number 5P41GM10339024) to P.A. at the Complex Carbohydrate Research Center, and the Cariplo Foundation [2016-0481], the Vontobel Foundation, and institutional funding of both the ETH Zurich and the University Hospital Balgrist to (J.G.S.).

## Supplementary Figures

**Figure S1.**
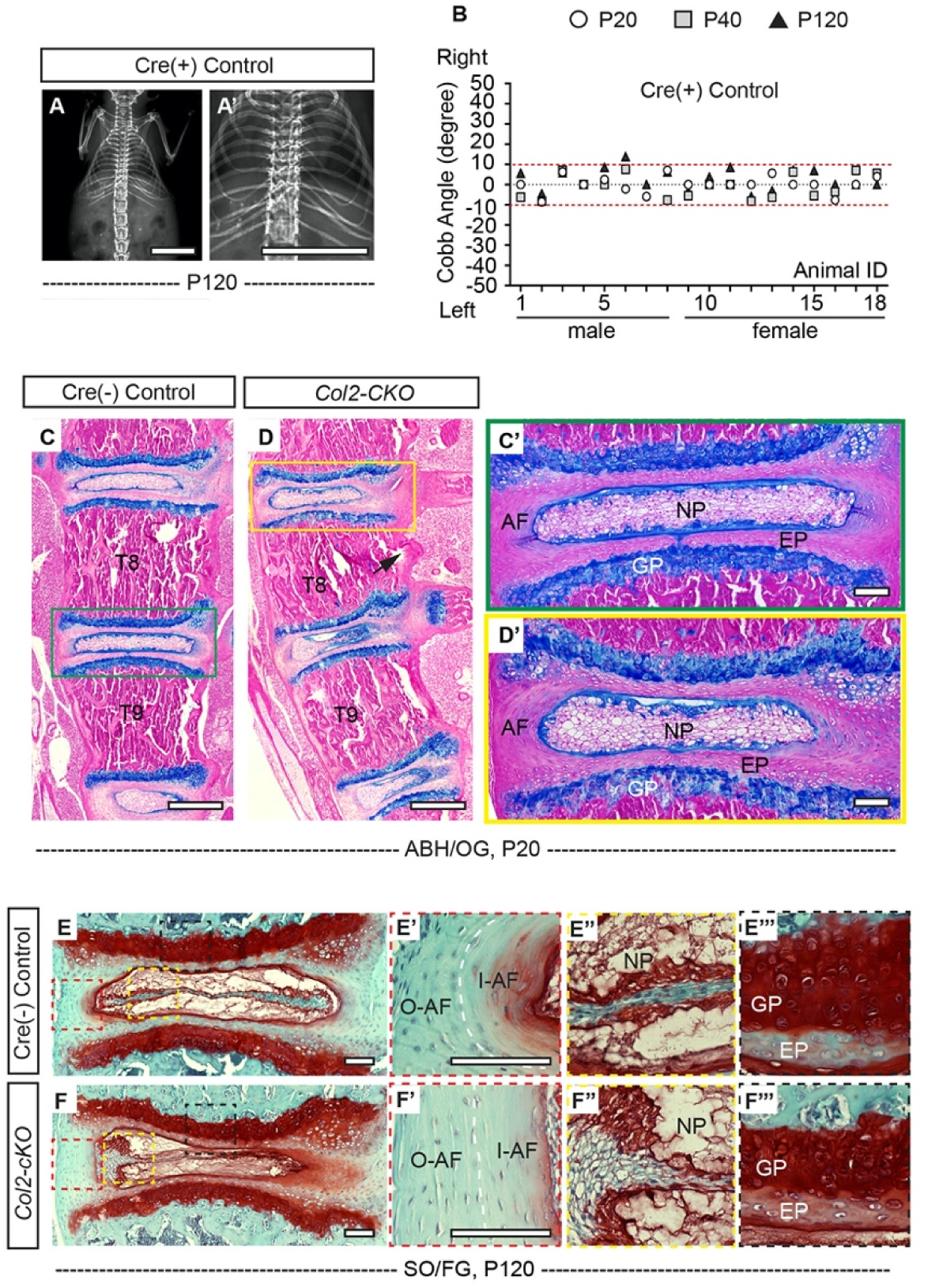
Loss of *Adgrg6* in osteochondral progenitor cells leads to alternations in spinal elements. **(A, A’)** X-ray analysis of a representative Cre (+) control mouse at P120. **(B)** Cobb angle values for all the Cre (+) control mice as shown in Fig. 1I. Thresholds of scoliosis (Cobb angle >10 degree) are indicated with two red dot lines. Only one Cre (+) control mice exhibited scoliosis at P120 (1/18). **(C-D’)** Representative thoracic spine (T8-T9) and IVD tissues of Cre (-) control and *Col2-cKO* mutant mice stained with Alcian Blue Hematoxylin/Orange G (ABH/OG) at P20. The mutant IVD outside of the spine curve (D’) was comparable with the control IVD (C’). Black arrow indicates the cortical bone bulged to the concave side of the curvature in the apical region of the mutant spine (D). *n*=3 for each group. **(E-F’’’)** Representative IVD tissues of Cre (-) control and *Col2-cKO* mice stained with Safranin-O/Fast green (SO/FG) at P120. We can observe exacerbated phenotype of wedged IVD (F), more vertical lamellae in the I-AF and O-AF (white dash line, F’), sifted nucleus pulposus (F”), and disorganized EP and GP (F’’’) in the mutant mice. *n*=3 for each group. Scale bars: 10mm in A, A’; 100μm in C-D’ and E-F’. *AF- annulus fibrosis, I- AF- inner annulus fibrosis, O-AF-outer annulus fibrosis, EP- endplate, GP- growth plate, and NP- nucleus pulposus*.

**Figure S2.**
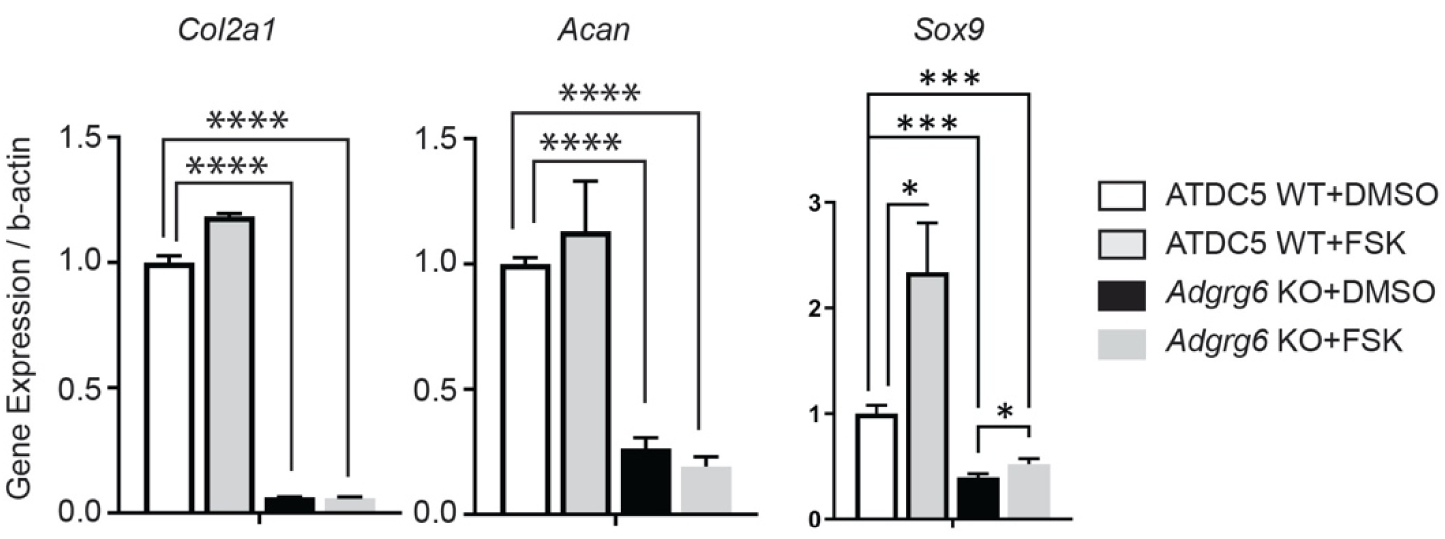
Forskolin treatment can partially rescue the dysregulation of *Sox9* expression but not *Col2a1* and *Acan* expression in *Adgrg6* KO cells. Real-time RT-PCR analyses of *Col2a1*,*Acan* and *Sox9* in wild type ATDC5 cells (ATDC5 WT) and *Adgrg6* knockout ATDC5 cells (*Adgrg6* KO), cultured with 2µM of Forskolin (FSK) or DMSO control for 7 days with maturation medium. The decreased expression of *Col2a1* and *Acan* in *Adgrg6* KO cells was not rescued by FSK treatment. The decreased expression of *Sox9* was partially rescued by FSK treatment. *n*=3 biological replicates, and the representative result is shown. Bars are plotted with mean and SD. ***: *p*<0.005, ****: *p*<0.001, One-Way ANOVA followed by Tukey test.

**Figure S3.**
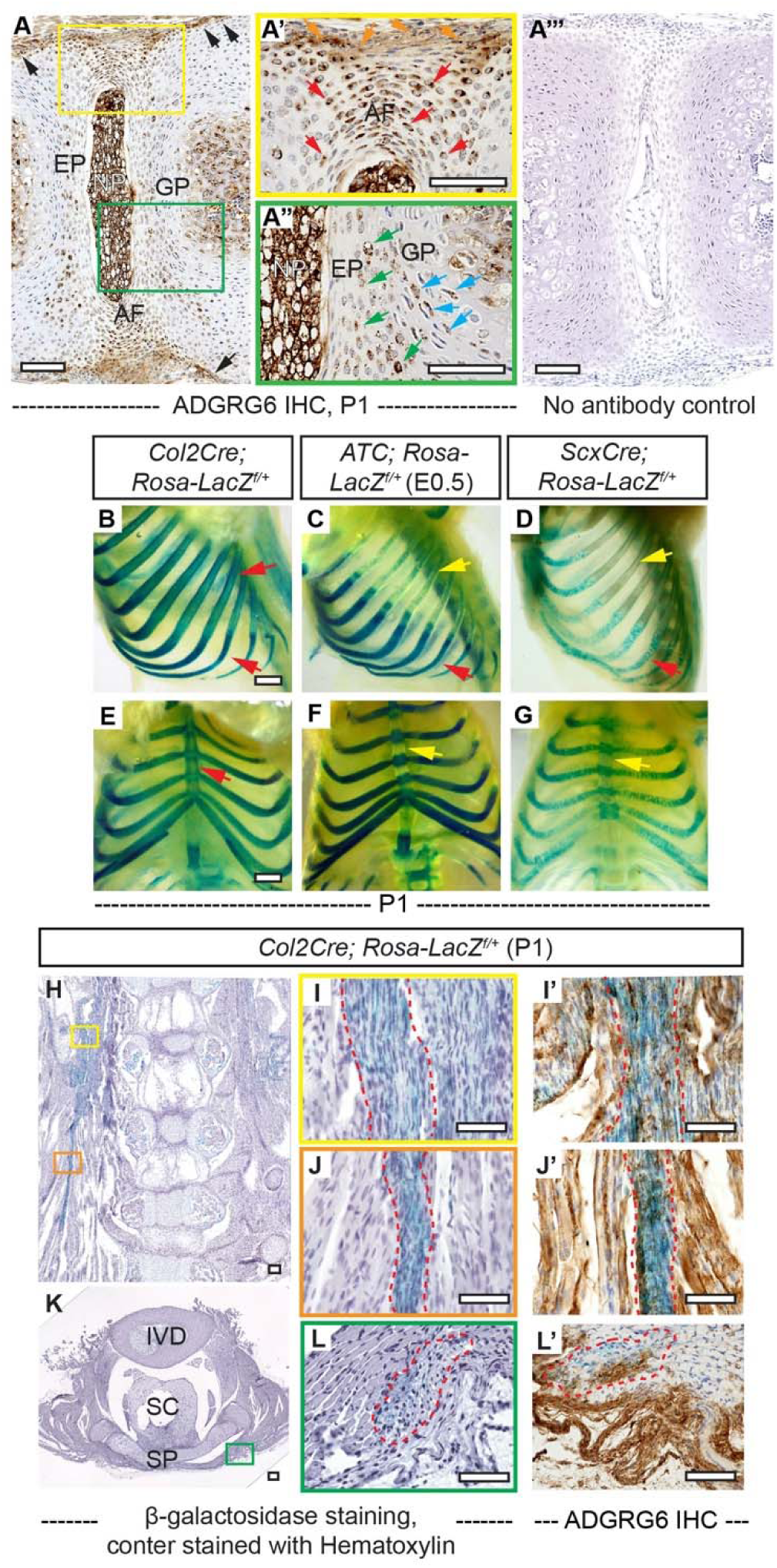
IHC analyses of ADGRG6 and β-galactosidase staining of *Rosa-LacZ* reporter mice recombined with different Cre strains. **(A-A’’’)** IHC analyses of ADGRG6 on thoracic spine sections of wild type mice at P1. ADGRG6 is mainly expressed in AF (red arrows, A’), NP (A”), EP (green arrows, A”), and GP (blue arrows, A”). Note that ADGRG6 also expressed in the outmost AF (orange arrows, A’) and the periosteum (black arrows, A). No antibody control staining is shown in A’’’. *n*=3 mice. **(B-G)** Whole mount β-galactosidase staining of *Rosa-LacZ* reporter mice at P1. Sagittal and ventral views of *Col2Cre; Rosa-LacZ^f/+^* mouse (B, E) and *ATC; Rosa-LacZ^f/+^* mouse (C, F, induced from E0.5) show robust recombination within cartilaginous tissues of the rib cage (red arrows, B and C). Sagittal and ventral views of *ScxCre; Rosa-LacZ^f/+^* mouse (D, G) only show some recombination signals within cartilaginous portions of the rib cage (red arrow, D). Note that *Col2Cre* targets both bony and cartilaginous portions of the ribs (red arrows, B), while *ATC* and *ScxCre* do not target the bony ribs (yellow arrows, C and D). In addition, the sternum is only targeted by *Col2Cre* (red arrow, E), but not *ATC* or *ScxCre* (yellow arrows, F and G). **(H-L’)** Coronal (H-J’) and transverse (K-L’) sections of the *Col2Cre; Rosa-LacZ^f/+^* mouse spine as shown in Fig. 3N. Blue signal represents recombination within the ligamentous tissues, which is also outlined with red dot lines in (I-L’). IHC analyses of ADGRG6 performed on adjacent sections of I, J, and L are shown in I’, J’ and L’. Most cells that are targeted by *Col2Cre* (blue signal) also express ADGRG6 (brown signal). *n*=3 mice in each group. Scale bars: 100μm in A-A’’’ and H-L’; 1mm in B, E. *AF- annulus fibrosis, EP- endplate, GP- growth plate, NP- nucleus pulposus, IVD- intervertebral disc, SC-spinal cord, and SP- spinous process*.

**Figure S4.**
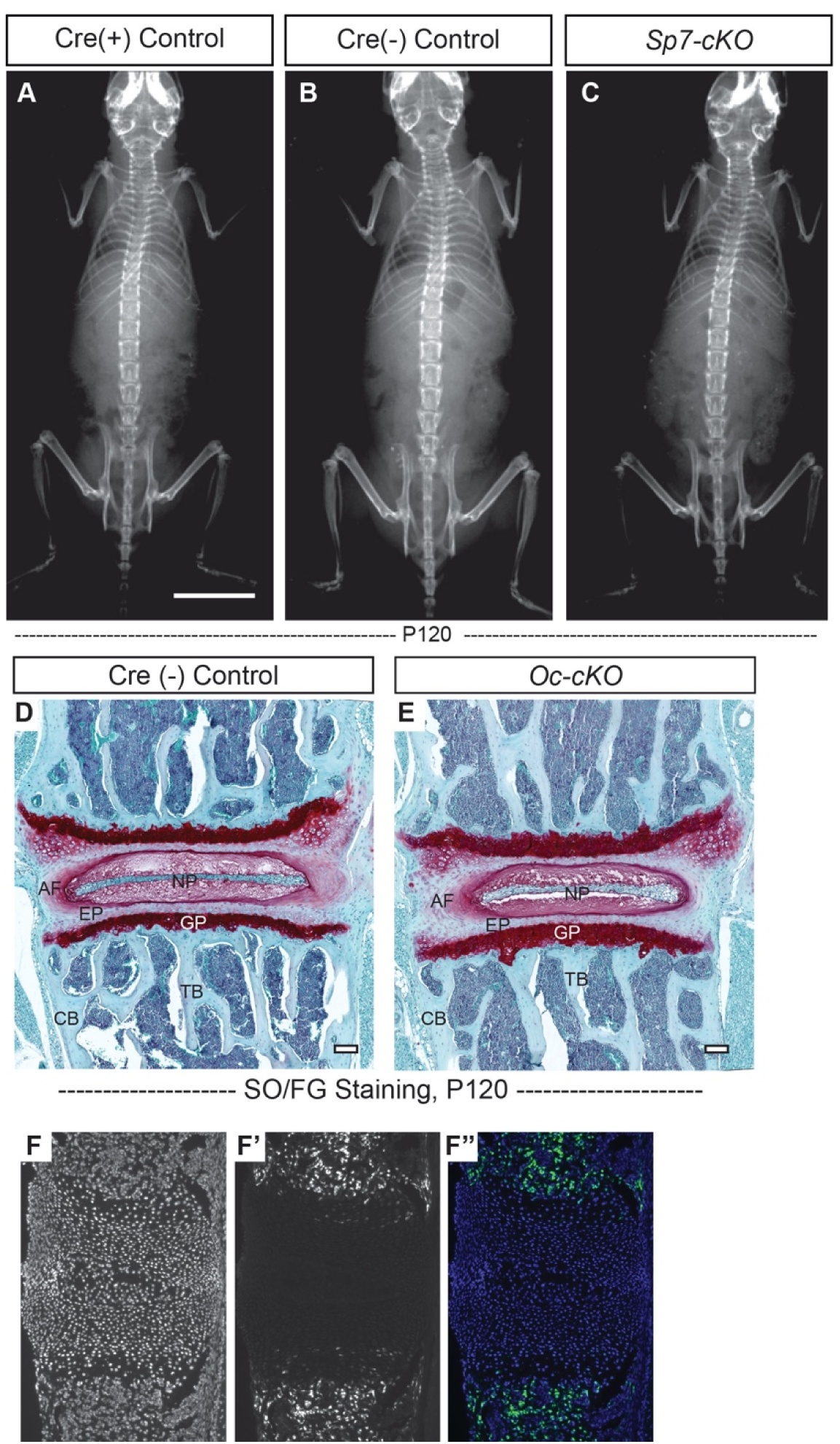
Loss of *Adgrg6* in osteoblast lineages leads to no scoliosis or spinal deformity. **(A-C)** Representative X-ray images of Cre (+) control (A), Cre (-) control (B), and *Sp7-cKO* mutant (C) mice at P120. *n*= 6, 5, 9 for Cre (+) control mice, Cre (-) control mice, and *Sp7-cKO* mice, respectively. **(D, E)** Representative IVD and vertebral body sections of Cre (-) control and *Oc-cKO* mutant mice stained with Safranin-O/Fast green (SO/FG) at P120. **(F-F’’)** Midline sectioned P1 mouse spine showing *Sp7-Cre-GFP* expression specific to the vertebral body (F’, F’’), counterstained with DAPI (F). *n*=3 mice for each group. Scale bars: 10mm in A-C; 100μm in D, E. *AF- annulus fibrosis, EP- endplate, GP- growth plate, NP- nucleus pulposus, TB-trabecular bone, and CB-cortical bone*.

**Figure S5.**
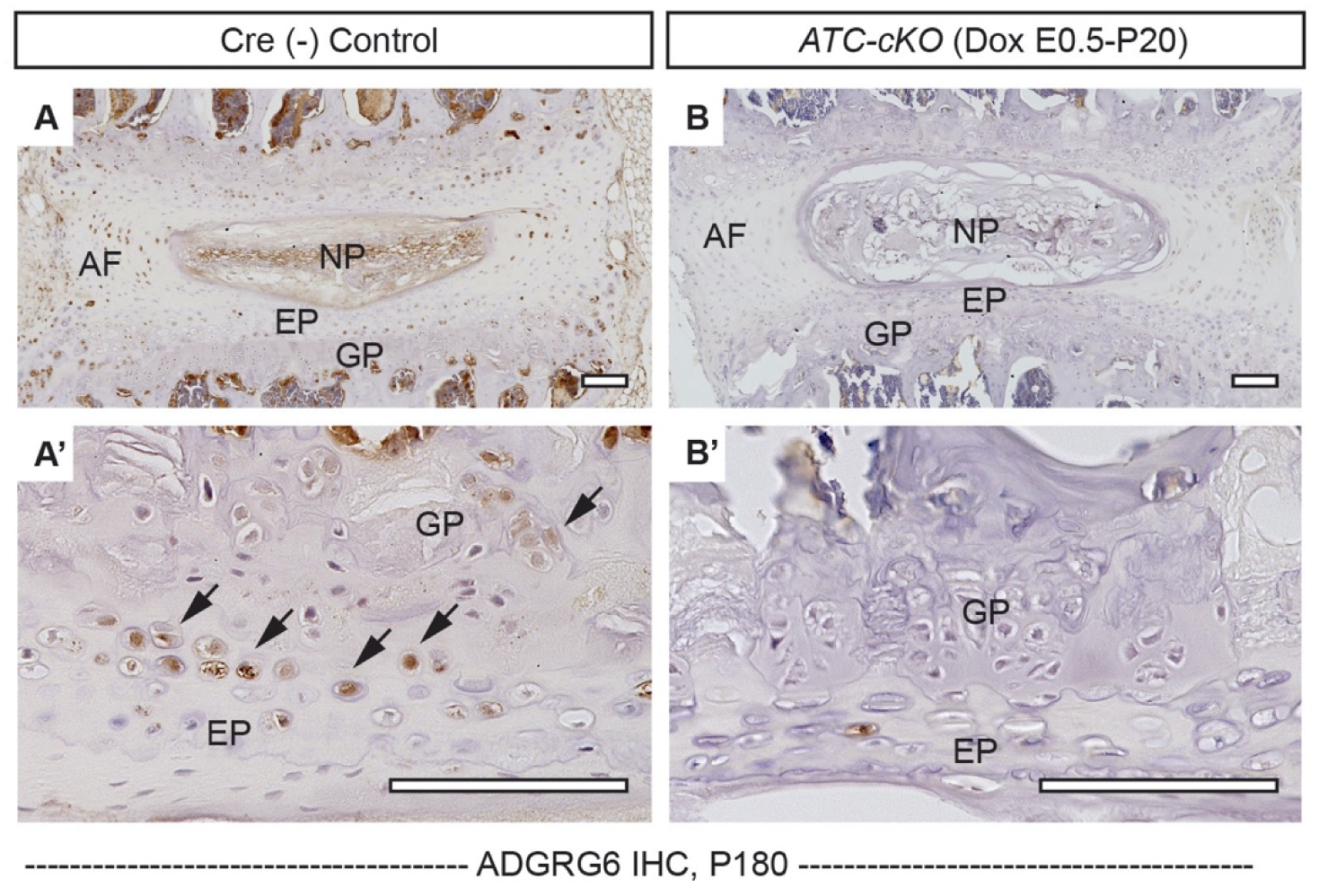
ADGRG6 expression is dramatically reduced in *ATC-cKO* mouse. Representative IHC analyses of ADGRG6 on spine sections of Cre (-) control mouse (A, A’) and *ATC-cKO* mouse (induced from E0.5-P20) (B, B’). ADGRG6 expression is dramatically reduced in *ATC-cKO* mouse. *n*=3 mice for each group. Scale bars: 100 µm. *AF- annulus fibrosis, EP- endplate, GP- growth plate, NP- nucleus pulposus*.

**Figure S6.**
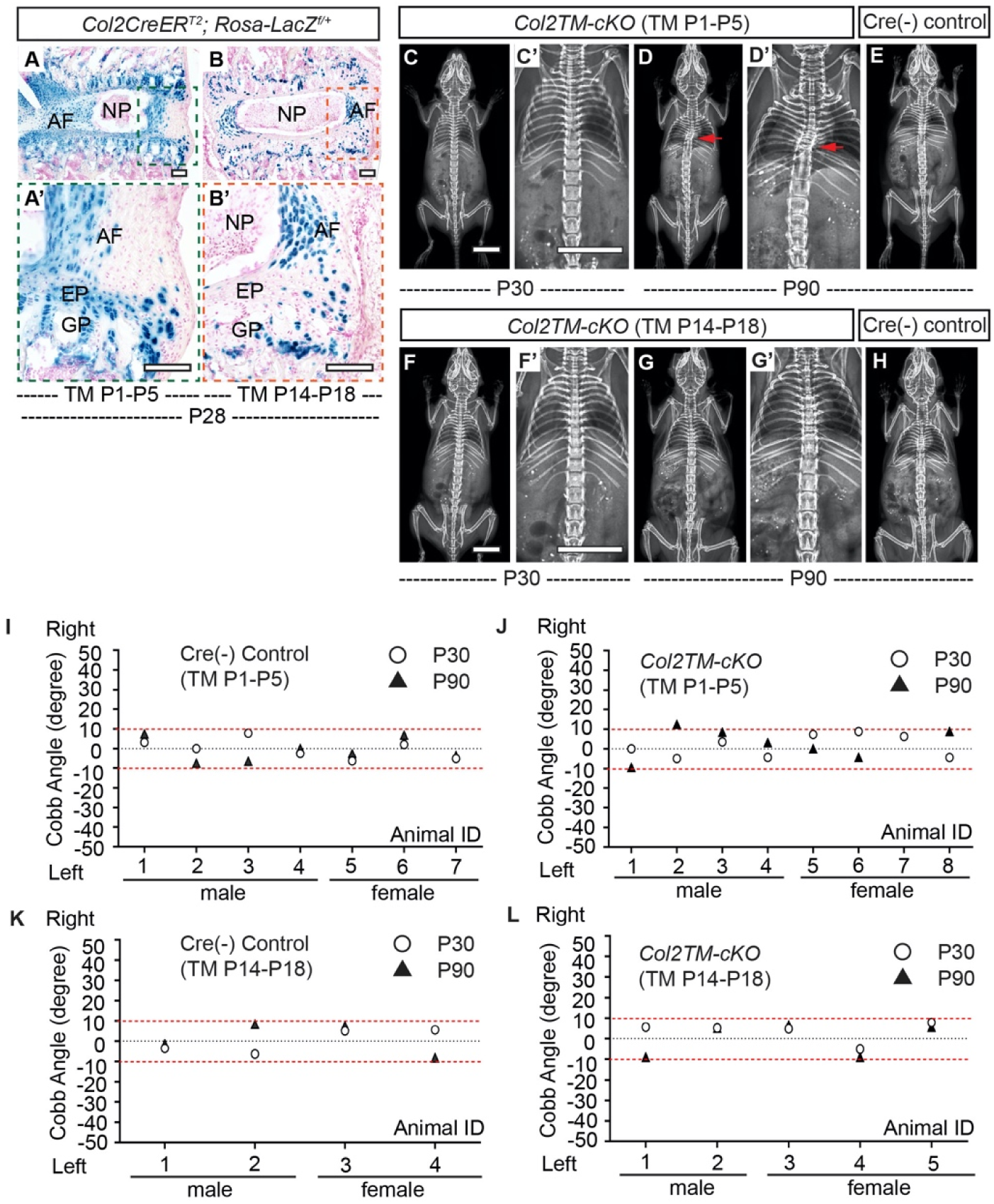
Loss of *Adgrg6* in cartilaginous tissues leads to low penetrance of scoliosis. **(A-B’)** β-galactosidase staining of *Col2CreER^T2^; Rosa-LacZ^f/+^* spine sections that were inducted from P1-P5 (A, A’) or P14-P18 (B, B’) at P28. Note that *Col2CreER^T2^* targets EP, GP and inner AF, but does not target NP or outer AF with neither induction strategy (A’, B’). **(C-H)** Representative X-ray images of *Col2TM-cKO* mutant mice (TM induced from P1-P5 or from P14-P18) at P30 (C, C’ or F, F’, respectively) and P90 (D, D’ or G, G’, respectively). Corresponding Cre (-) controls at P90 are shown in (E) and (H). Scoliosis is indicated with red arrows in (D, D’). **(I-L)** Longitudinal analyses of Cobb angle values of Cre (-) control mice and *Col2TM-cKO* mice at P30 and P90 with two induction strategies. For TM induced from P1-P5, *n*= 7 mice for Cre (-) controls at both P30 and P90; *n*= 8 and 7 mice for *Col2TM-cKO* mice at P30 and P90, respectively. For TM induction from P14-P18, *n*=4 mice for Cre (-) controls at both P30 and P90; *n*=5 for *Col2TM-cKO* mice at both P30 and P90. Thresholds of scoliosis (Cobb angle >10 degree) are indicated with two red dot lines. Scale bars: 100μm in A-B’; 10mm in C, C’ and F, F’. *AF- annulus fibrosis, EP-endplate, GP- growth plate, and NP- nucleus pulposus*.

**Figure S7.**
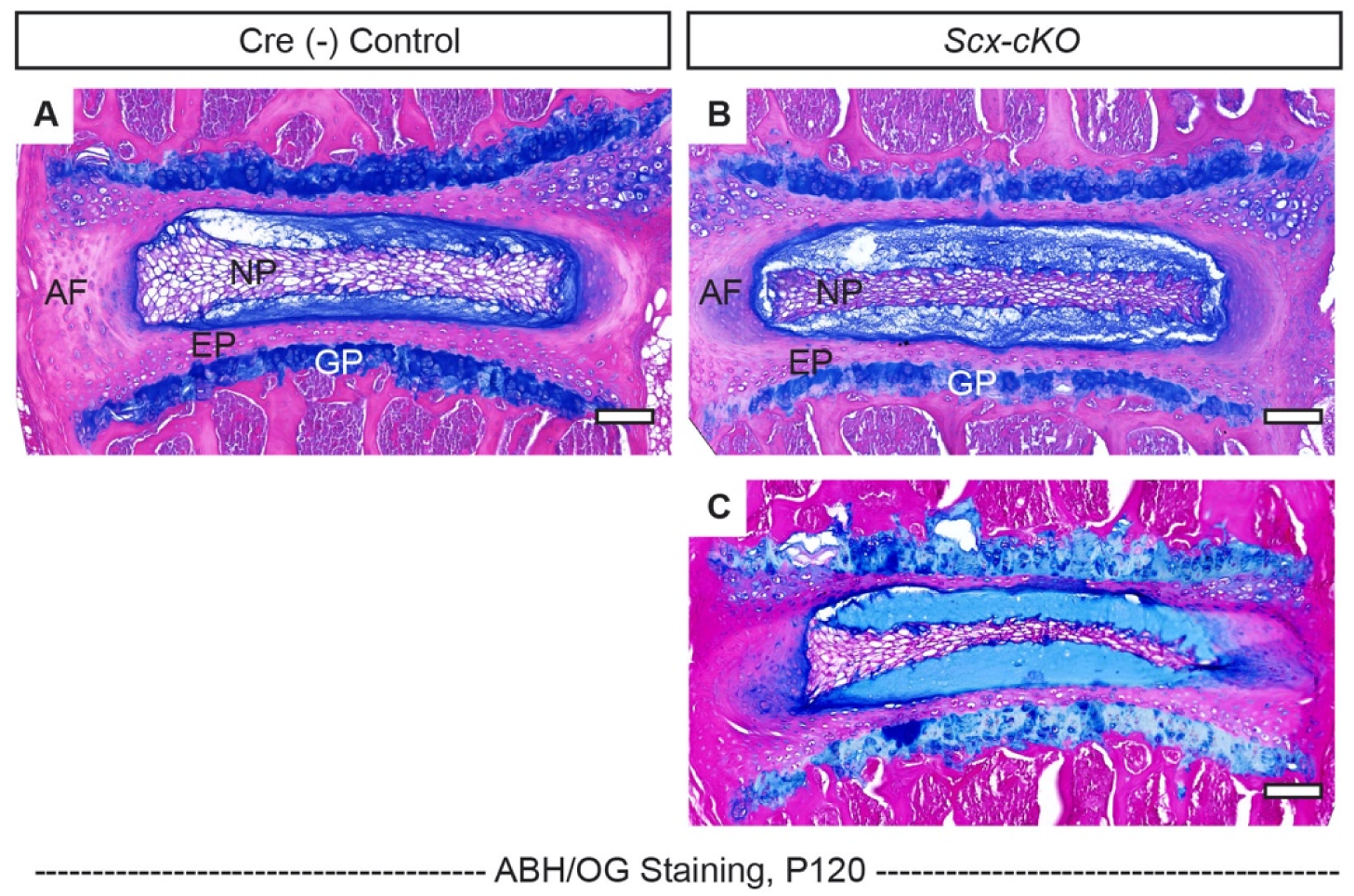
*Scx-cKO* mice display mildly wedged IVDs within the curve. **(A-C)** Representative IVD tissues of Cre (-) control (A) and *Scx-cKO* mutant mice (B, C) stained with Safranin-O/Fast green (SO/FG) at P120. IVD outside of the curve and within the curve are shown in (B) and (C), respectively. *n*=3 mice for each group. Scale bars: 100μm. *AF- annulus fibrosis, EP- endplate, GP- growth plate, and NP- nucleus pulposus*.

**Table S1:** Chondroitin sulfate digestion profile and total hyaluronan content in Cre (-) control and *Col2-cKO* mutant mice at P20

|  | Control-1 |  | Control-2 |  | Control-3 |  | Mutant-1 |  | Mutant-2 |  |
| --- | --- | --- | --- | --- | --- | --- | --- | --- | --- | --- |
| **CS** |  |  |  |  |  |  |  |  |  |  |
| D0a0 | 3.88 | 4.0 | 6.10 | 4.6 | 7.35 | 3.8 | 6.30 | 5.4 | 4.45 | 5.1 |
| D0a6 | 9.34 | 9.5 | 11.31 | 8.5 | 15.96 | 8.3 | 9.56 | 8.1 | 6.99 | 8.0 |
| D0a4 | 83.25 | 85.1 | 113.92 | 85.5 | 167.09 | 87.0 | 100.49 | 85.4 | 75.05 | 86.1 |
| D2a0 | ND |  | ND |  | ND |  | ND |  | ND |  |
| D2a6 | ND |  | ND |  | ND |  | ND |  | ND |  |
| D0a10 | 0.25 | 0.3 | 0.63 | 0.5 | 0.34 | 0.2 | 0.30 | 0.3 | 0.14 | 0.2 |
| D2a4 | 1.12 | 1.1 | 1.30 | 1.0 | 1.19 | 0.6 | 1.09 | 0.9 | 0.59 | 0.7 |
| D2a12 | ND |  | ND |  | ND |  | ND |  | ND |  |
| Total CS | 97.84 | 100 | 133.26 | 100 | 191.92 | 100 | 117.73 | 100 | 87.22 | 100 |
| Total HA | 4.09 |  | 3.41 |  | 1.44 |  | 5.41 |  | 2.05 |  |
| Total GAG | 101.93 |  | 136.67 |  | 193.36 |  | 123.14 |  | 89.27 |  |

(µg estimated for total reaction volume; *w/w percentage*)
CS: chondroitin sulfate; HA: hyaluronan; GAG: glycosaminoglycan
ND = Not Detected

## Notes

### Competing Interest Statement

The authors have declared no competing interest.

